# Neural and computational mechanisms of effort under the pressure of a deadline

**DOI:** 10.1101/2024.04.17.589910

**Authors:** M. Andrea Pisauro, Daniele Pollicino, Lucy Fisher, Matthew A. J. Apps

## Abstract

Deadlines fundamentally shape the motivation for effort. Research examining effort-based choices finds high effort is an avoided cost. However, this work overlooks the fact that effort can be valuable when it makes progress on long-term goals before deadlines. We test a new framework where motivation depends on deadline pressure (work remaining / time remaining). Across three studies we use computational modelling on novel tasks examining effort-based decisions when effort makes progress on goals with deadlines. In support of hypotheses, deadline pressure significantly impacts decision-making, shifting people from avoiding effort, to seeking and valuing it. Using ultra-high-field fMRI, we show that functionally connected putamen and midcingulate cortex (MCC) sub-regions process and update estimates of deadline pressure, with distinct anterior cingulate and putamen sub-regions processing the costs or added value of effort. We show the neurocomputational mechanisms for how deadline pressure shapes motivation, and that keep us ‘on track’ for our goals.

## Introduction

Deadlines – a point in time by which a long-term goal must be completed – are a fundamental component of modern societies across education, employment, and sporting endeavours and also of the natural world. When the squirrel decides to exert themselves to find and bury more food before winter^1^, the athlete trains hard for a marathon, or the academic chooses to put all their effort into a grant application^2–5,6^; Our goals are often achieved only if high amounts of effort are chosen over time before a deadline. Yet, while there is research separately examining how goals^7,8^, task progress^9^ and effort^10–13^ are processed in the brain, none has put these multiple components that have major impacts on motivation together. Thus, fundamental questions about the computational and neural mechanisms underlying how the brain makes decisions to exert effort to progress on goals when under the pressure of a deadline are unanswered.

The willingness to exert effort is vital for foraging animals^14–17^ and human health^1,6^. Yet, research examining the mechanisms of effort-based decision-making has used tasks in which behaviours conform to a ‘law of less effort’^18^. In such tasks effort is avoided unless associated with large immediate extrinsic rewards, indicating it is treated as a cost to be avoided, and only chosen if sufficiently incentivised ^10,16,19–21^. In contrast, psychological theories have proposed that effort can also be valued, leading some to suggest effort is paradoxical^22^ being both an avoided cost and a sought benefit. Anecdotally people undertake highly effortful behaviours associated with little immediate extrinsic reward with regularity, such as training for a marathon or climbing a mountain suggesting effort is not always a cost. However, while there is evidence animals can change their preferences for how willing they are to exert effort over time^2,23–25^, and people self-report liking effortful behaviours^22,26^, there is little empirical evidence that people actually seek and value a higher effort behaviour, nor an understanding of the neural mechanisms underlying this.

Studies examining the neural and computational mechanisms of effort also typically assume that effort is a cost that discounts the value of rewards^10,21,27^. Models of effort-discounting have provided a robust account of this effort aversion, and when combined with neuroscience methods have revealed several frontal and basal-ganglia circuits linked to decisions of whether to exert effort^10,11,21,23,28^. Lesions to these regions in humans, and homologous areas in animals models, reduce the willingness to exert effort^16,21,29^. Portions of the sulcus of the midcingulate cortex (MCC), typically labelled dorsal anterior cingulate cortex (dACC), and putamen are consistently shown to signal the costs of effort and the subjective value of rewards discounted by effort^10,23,28,30^. Notably, individual differences in responses to efforts’ costs in MCC and putamen are associated with variability in effort aversion^28^. In addition, ACC, ventro-medial prefrontal cortex (VMPFC), anterior insula (AI), and ventral striatum (VS) have all been linked to processing the rewards and subjective value of rewards discounted by efforts costs^11,23,28,31,32^.

Strikingly, variability in neural responses in sub-regions of the MCC and ACC have also been associated with changes in activity when one progresses through a sequential task^9,12,33^, as levels of fatigue fluctuate impacting effort-based decisions,^23^ with motivating extended behaviours^33^, and with shifting preferences for sequential risky choices when under pressure to complete a goal^34^. Putting these findings together might suggest these regions could perform functions that shift the valuation of effort, and thus decisions of whether to exert effort for reward, when making progress towards goals. However, there are no studies that provide any evidence of this, nor a formalised account of the computations that would drive such a shift.

Here, we offer a normative explanation and formal computational model for how the pressure to reach longer-term goals before deadlines can shift effort from a cost to something positively valued. We propose that longer-term goals are only reached if effort is exerted. Typically, greater effort leads to more progress being made towards completing a goal. When there is no deadline, progress can be made slowly, and high effort is not necessary. However, when there is a deadline, all the progress necessary to complete the goal needs to be made by a fixed point in time, and therefore small amounts of effort may be insufficient to reach the goal in time. That is, if you are close to a deadline in time and have significant progress required to complete a goal, effort has high value as it is a necessity in order to reach the goal before the deadline^2^. Thus, there is a high pressure to choose it. Alternatively, if you are close to a deadline but have nearly completed the goal, pressure is low, and effort will not add value. Thus ‘deadline pressure’, defined as the amount of work still required to complete a goal divided by the time left to complete it, should guide decisions of when to exert effort to complete goals. We propose that this deadline pressure switches the valence of effort within decision-making computations, leading effort to be valued. Thus, the more deadline pressure an individual is under, the more they will value effort. Moreover, sub-regions of the MCC, ACC, VMPFC, AI and VS previously implicated in effort and goal processing will process the range of computations underlying this deadline pressure, and how effort can be at times an avoided cost but at others valued and sought.

To test these hypotheses, we developed a novel effort-based decision-making task. In this task, people have to exert effort in order to make progress towards goals. The more effort they exert, the more progress they make towards the goal. However, the goals must be completed in 8 trials – creating a deadline – in order to increase a bonus payment for participation. On each trial, they chose between lower or higher effort options, that both vary in the amount of effort (grip force or button presses) required.

Across three studies using computational modelling (Study 1: ultra-high-field fMRI; Study 2: lab behaviour only; Study 3: online) we show that people’s choices are strongly influenced by deadline pressure. When there is more pressure people choose the higher effort option more frequently with modelling revealing that this is because deadline pressure causes the amount of effort to add to the value of offered rewards, rather than being a cost. We show that that individual differences in the sensitivity to pressure is associated both with being more proactive in the task (choosing the higher effort earlier to reach a goal) and with self-reported daily levels of fatigue. We found that there was a spectrum of responses across functionally connected sub-regions of the MCC, ACC and putamen that signalled features of the computational model including deadline pressure and the subjective added value of effort, as well as processing the chosen amount of effort when it was being treated as valuable, with separate regions signalling it when treated as a cost. These findings highlight that the pressure caused by deadlines switches the neurocomputational mechanisms of effort, leading to the processing of effort as valuable and worth seeking, rather than a cost to be avoided.

## Results

To test the hypothesis that the pressure of a deadline switches effort from an avoided cost to something valuable, and the neural mechanisms underlying how to choose whether to exert effort to reach goals before a deadline, across three studies participants performed novel decision-making tasks. In these tasks the exertion of physical effort made progress towards a goal to be reached within a fixed deadline after 8 trials (**Fig. 1**). Like many goals in real-life, if they make all the progress required to complete a goal, they can receive all of the reward, if they do not, they will not obtain rewards (e.g., if a student fails to finish an essay, they will not get a positive grade). In each trial, subjects were offered a choice between two options that varied in terms of the levels of effort required (20 – 70% of their maximum voluntary contraction [MVC] / grip strength) and the levels of reward on offer, represented as coins (**Fig.1a**). The choice was always a trade-off between collecting more coins for less effort or exerting more effort for less coins (i.e. the opposite of typical effort discounting tasks). However, as more effort led to greater amounts of progress towards the goal, participants needed to choose the higher effort, lower reward option multiple times in order to reach the goal before the deadline. After each decision, they were asked to exert the effort associated with their choice by squeezing a hand grip at the chosen level for 1 out of 3s. If they reached the required level, they progressed towards the goal by an amount proportional to the effort exerted. If they failed, they did not progress nor collected any coins in that trial (**Fig. 1**). In study 1 participants performed the task inside a 7T fMRI scanner with the effort required being grip force. Study 2 was identical but performed in the lab, aiming to replicate the main behavioural results. In study 3 participants performed a task online with effort manipulated in terms of button presses rather than grip force (a proportion – varying from 50% to 95% – of the maximum voluntary number of clicks) with the aim to examine whether sensitivity to deadline pressure was associated with individual differences in psychological traits.

**Figure 1.**
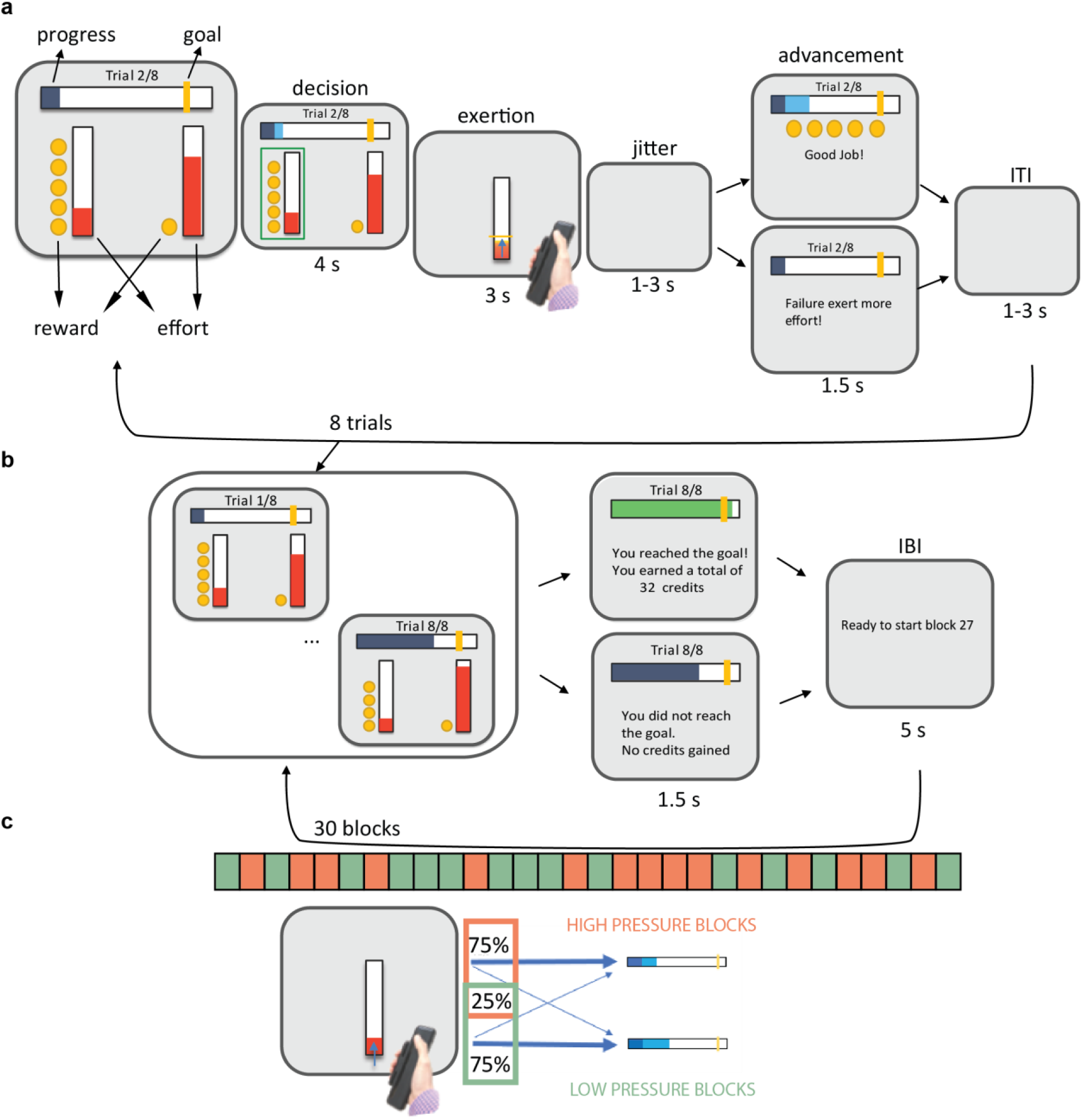
**a** Task design and trial structure. Participants need to reach a goal by exerting physical effort to complete goals before a deadline after 8 trials. Exerting more effort made more progress towards the goal. The goal is represented by a yellow line on the right-hand side of the horizontal bar (first panel). Participants’ progress towards this bar is indicated by the area filled in blue. In each trial subjects are offered a choice between two different levels of effort, represented by the red vertical bars with more physical effort a higher red bar, in exchange for two different amounts of reward, represented by the yellow coins. The choice was always a trade-off between collecting more coins or exerting more effort in order to advance towards the goal. After each decision indicated by the green line (2nd panel), subjects are asked to exert the effort associated with their choice by squeezing a hand grip (3rd panel). If they have reached the required level they advance towards the goal by an amount proportional to the effort exerted (4^th^ panel, top). If they fail, they don’t advance (4^th^ panel, bottom). **b**. Subjects were presented with 30 goals (30 blocks of 8 trials). If they do reach the goal, they receive all the reward accumulated throughout the block. If they don’t reach the goal, they earn nothing in that block. **c.** To increase the variability in how much pressure there was to reach goals, in each trial effort was mapped onto advancement towards the goal in a probabilistic fashion. In study 1 and 2, half of the blocks were “low pressure” blocks where there was a 75% chance of an enhancement in advancement with respect to effort exerted and a 25% chance of reduction. In the other half of blocks, in the “high pressure blocks”, the probabilities were reversed, with a 75% chance of reduction and 25% of enhancement of the mapping of effort into advancement. In study 3 the mapping of effort into advancement was kept fixed.

## Deadline pressure leads to seeking effort

Our hypothesis was that people would succeed at completing goals before deadlines by computing a “deadline pressure”, calculated for each trial as the amount of work left to be completed to reach the goal divided by the time (number of trials) left to complete it. This deadline pressure would then shift the value of effort to positive in choices, rather than being treated as a cost. Participants were indeed successful at completing goals in the task (Study 1: (65.8 ± 1.6%) average goals completion rate, collecting 698 ± 37 coins, **Fig.2c/Suppl. Fig. 1**; study 2: (70.4± 1.5%) goals completion rate, collecting 924 ± 74 coins, **Suppl. Fig. 1**; study 3: (62.1 ± 1.6%) goal completion rate, collecting 810 ± 96 coins; **Suppl. Fig. 1).**

While successfully reaching the goals indicated people were choosing higher efforts often enough to succeed, we next wanted to examine whether deadline pressure was driving such decisions. Using a logistic regression separately including the effort required and the reward on offer of both options, and deadline pressure (distance between the progress made and the goal / trials left to complete it; **Fig.2d**) as predictors of higher effort choices, we found highly significant effects of pressure across studies (Study 1: *β* = 26.9±1.2, *t* = 23.031; *p*<0.001; study 2: *β* = 19.1 ±1.5, *t* = 12.35; *p*<0.001; study 3: *β* = 13.5±0.9, *t* = 14.7; *p*<0.001; **Suppl. Fig. 3).** This logistic regression model structure was selected through model comparison (BIC) and outperformed those in which the advancement towards the goal, or time left at trial *t,* were included as separate variables, indicating that an integration of these variables into deadline pressure was a more likely computation guiding choices.

To further test whether participants responded adaptively as pressure varied, in the first two studies the mapping between the advancement towards the goal, and the effort exerted, was probabilistic (**Fig.1c**). Although participants were aware that the same amount of effort would not always lead to the same amount of progress towards the goal, unbeknownst to participants half of the blocks had lower pressure as on 75% of trials progress was easy (i.e. although there was still a direct mapping between effort exerted and progress, sometimes the amount of progress made by an effort would be higher) and on 25% it was hard (it required more effort), and half were higher pressure with the opposite proportion of hard and easy progress trials. This manipulation of effort in different blocks influenced participant choices, with people choosing the higher effort more often in high pressure than low pressure blocks (Study 1: *t* =11.48 p<0.001; **Fig.2a**; Study 2: *t* =3.33 p<0.005), and logistic regressions showing in study 1 a higher effect of effort on the higher effort option in high compared to low pressure trials (although this effect didn’t replicate in study 2; Study 1: *Δβ_high_effort_* = 0.25, *t* = 0.31; *p*<0.05; study 2: *Δβ_high_effort_* = –0.318, *t*=-0.32, *p* = 0.745). When advancement towards a goal was unexpectedly lower following an exerted effort participants responded by subsequently choosing the higher effort option more often (*t* = 6.99, *p*<0.001 for study 1 and *t*=7.26, *p*<0.001 for study 2). The block type (high/low pressure) significantly modulated the performance in the task, changing the success rate (by 31% in study 1, p < 0.001, and by 26% in study 2, p < 0.001; **Fig.2c and Suppl. Fig. 1**) and allowing participants to reach the goal earlier (by a mean of 0.6 trials in study 1, p < 0.001 and by 0.7 in study 2; p < 0.001, **Fig.2c and Suppl. Fig. 1**) in low pressure blocks compared to high pressure blocks. Average rewards collected did not differ significantly between high– and low-pressure blocks (**Suppl. Fig. 1**) nor did fatigue levels (measured by self-report between each block) differ significantly between the two kinds of blocks (*p* = 0.83 for study 1, *p* = 0.39 for study 2). Thus, participants were choosing the higher effort option more often when the offered effort was higher, as a function of the amount of pressure being caused by the deadline and how easy it was to progress on the task.

One possibility was that participants could be successful in this task, not by responding to pressure and valuing effort, but by simply choosing the more effortful option regardless of the levels of effort and reward on offer. Multiple analyses do not support that notion. Firstly, we found significant effects on choices of the offered amounts of effort and reward for both options, indicating that people were considering all aspects of the options presented to them (Effort study 1: *β_left_* = –4.59±0.51, *t* = –8.95; *p*<0.001; *β_right_* = 0.92±0.38, *t* = 2.41; *p*= 0.016; Reward study 1: *β_left_* = –0.217±0.016, *t* = –13.3; *p*<0.001; *β_right_* = 0.198±0.026, *t* = 7.7; *p*<0.001; Effort study 2: *β_left_* = –4.20±0.63, *t* = –6.69; *p*<0.001; *β_right_* = – 0.12±0.46, *t* = 4.64; *p* = 0.8; Reward study 2: *β_left_* = –0.238±0.020, *t* = –11.9; p<0.001; *β_right_* = 0.243±0.031, *t* = 7.8; *p*<0.001; Effort study 3: *β_left_* = –0.052±0.004, *t* = –13.7; *p*<0.001; *β_right_* = 0.094±0.003, *t* = 33.69; *p* <0.001; Reward study 3: *β_left_* = 0.341±0.013, *t* = 24.9; *p*<0.001; *β_right_* = –0.22±0.019, *t* = –11.5; *p*<0.001; **Suppl. Fig. 3**).

Secondly, as outlined above we showed that people sought the higher effort option more often the more it was higher in effort and also were more likely to choose the lower effort option when this lower effort was closer to the higher level (**Fig.2d; Suppl. Fig. 3**). This suggests that when people were trying to make progress towards the goal, they were more willing to choose either of the options the higher they were in effort, rather than the lower (this effect was replicated in study 3 but was only present for the lower effort in the smaller sample study 2). In simple terms, participants were more likely to pick options the more effortful they were, suggesting that they were prioritising making progress towards the goal and thus seeking high effort options that could do so. Taken together these results suggest that subjects experienced deadline pressure and this led them to prefer to exert effort rather than avoiding it.

Is it possible that the participants in these studies don’t show the typical effort discounting effect? To test this, the same participants performed a standard effort-based decision-making task (prior to the main task) where there was no goal. Here, a logistic regression revealed that the effect of the higher effort was aversive (p < 0.05), discouraging higher effort choices, whereas in the same regression in the main task the effect was positive (p < 0.05), with the presence of pressure inducing effort seeking behaviour (**Fig. 2e left panel and Suppl. Fig. 3**). Consistent with this, we ran a computational model (a parabolic effort-discounting model as commonly used for physical effort studies^10,11,35^) which revealed that discounting parameters (k) were positive for all subjects in the effort-discounting task fit, but when fitted to the main task with goals and deadlines, were all 0 (i.e. no discounting of rewards by effort) apart from one participant (**Fig.2e right panel**) supporting the notion that, under pressure, effort stops discounting reward and becomes valuable.

**Figure 2.**
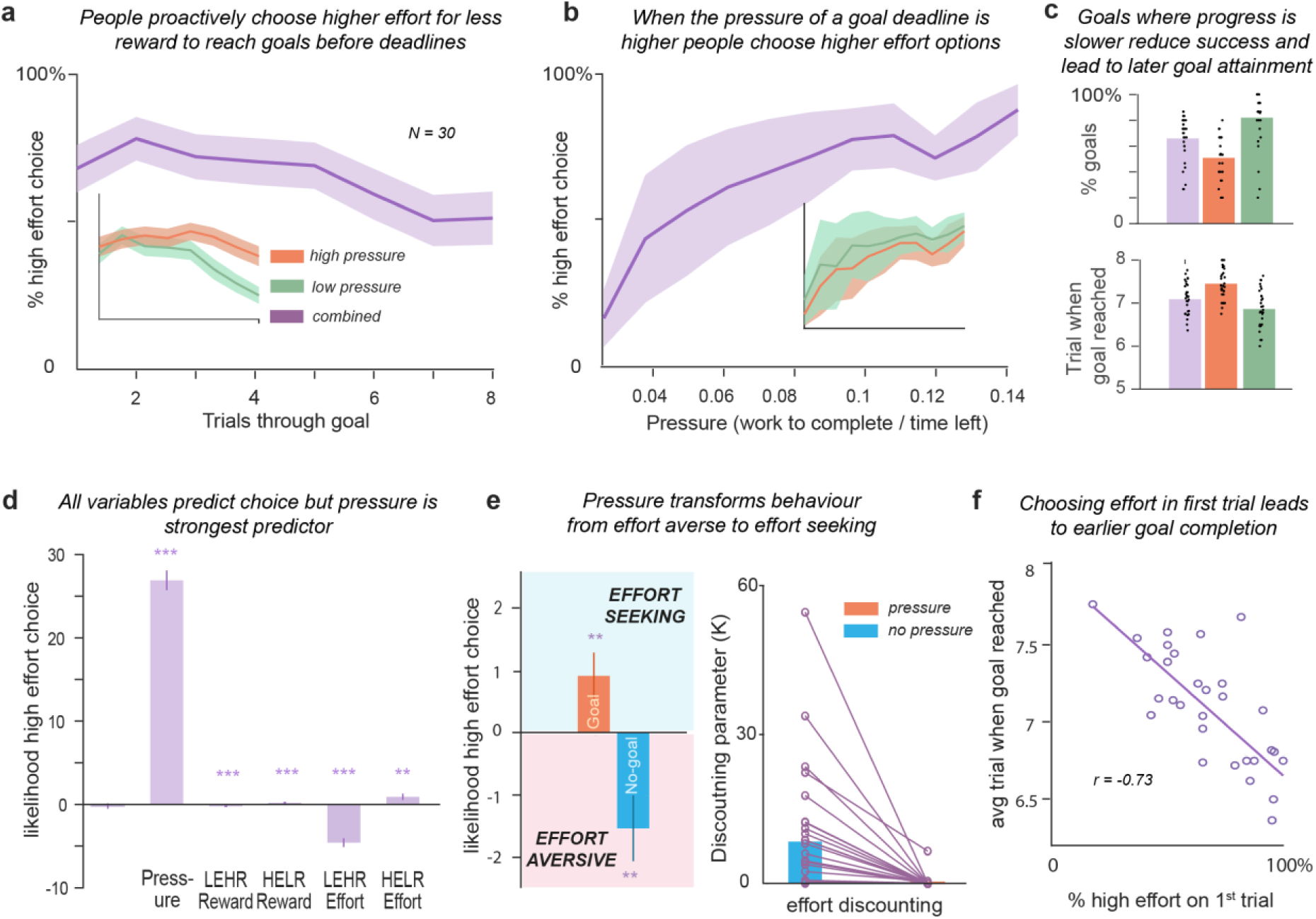
Behavioural results from study 1. **a**. Average (across subjects and blocks) percentage of higher effort choices across each trial number within blocks. In the inset, the same for high-pressure (orange) and low-pressure (green) blocks. **b**. Average (across subjects and blocks) percentage of higher effort choices across binned pressure values (10 bins). In the inset, the same for high-pressure (red) and low-pressure (green) blocks. **c**. Average percentage of goals reached across all (purple), low-pressure (green) and high-pressure blocks (orange; top panel) and average number of trials before deadline in which the goal was reached across all, high-pressure and low-pressure blocks (bottom panel). **d.** A logistic regression predicting choices in the main task as a function of pressure, the reward level of the lower effort option (LEHR), the reward of the higher effort option (HELR), the effort level of the lower effort option, and the effort level of higher effort option. **e.** Regression values (left panel) of the effort level of the higher effort option during the main task with a goal (orange) and of the higher effort option of the standard effort discounting task (blue). On the absence of pressure, participants were effort averse, but in its presence, they were effort seeking. Consistently, the discounting parameter (k) encoding how much effort was discounting rewards (methods) was positive when the model was fit to a standard effort discounting task without goals, where there was no pressure, but was near zero when the model was fit to the main task trials, in the presence of deadline pressure (right panel). **f.** The percentage of times participants choose the higher effort lower reward choice in the first trial (when pressure is the same for all participants) negatively correlated with the average trial at which participant reached the goal.

## Computational models reveal the added value of effort when under deadline pressure

To understand the computational mechanisms underlying choices, we developed multiple alternative models for people’s decisions of whether to exert effort in the task. Within this comparison we included models that assumed effort-discounting where effort is a cost that devalues rewards, models that followed a simple heuristic rule that would allow the reaching of goals but did not include effects of pressure (e.g. choose higher effort until goal is reached then choose low effort), models that included information about time left and advancement towards the goal separately and not integrated into a pressure term, and several models of deadline pressure which characterised how pressure might modulate different aspects of how choices were made in the task in different ways (Methods; **Fig. 3**).

**Figure 3.**
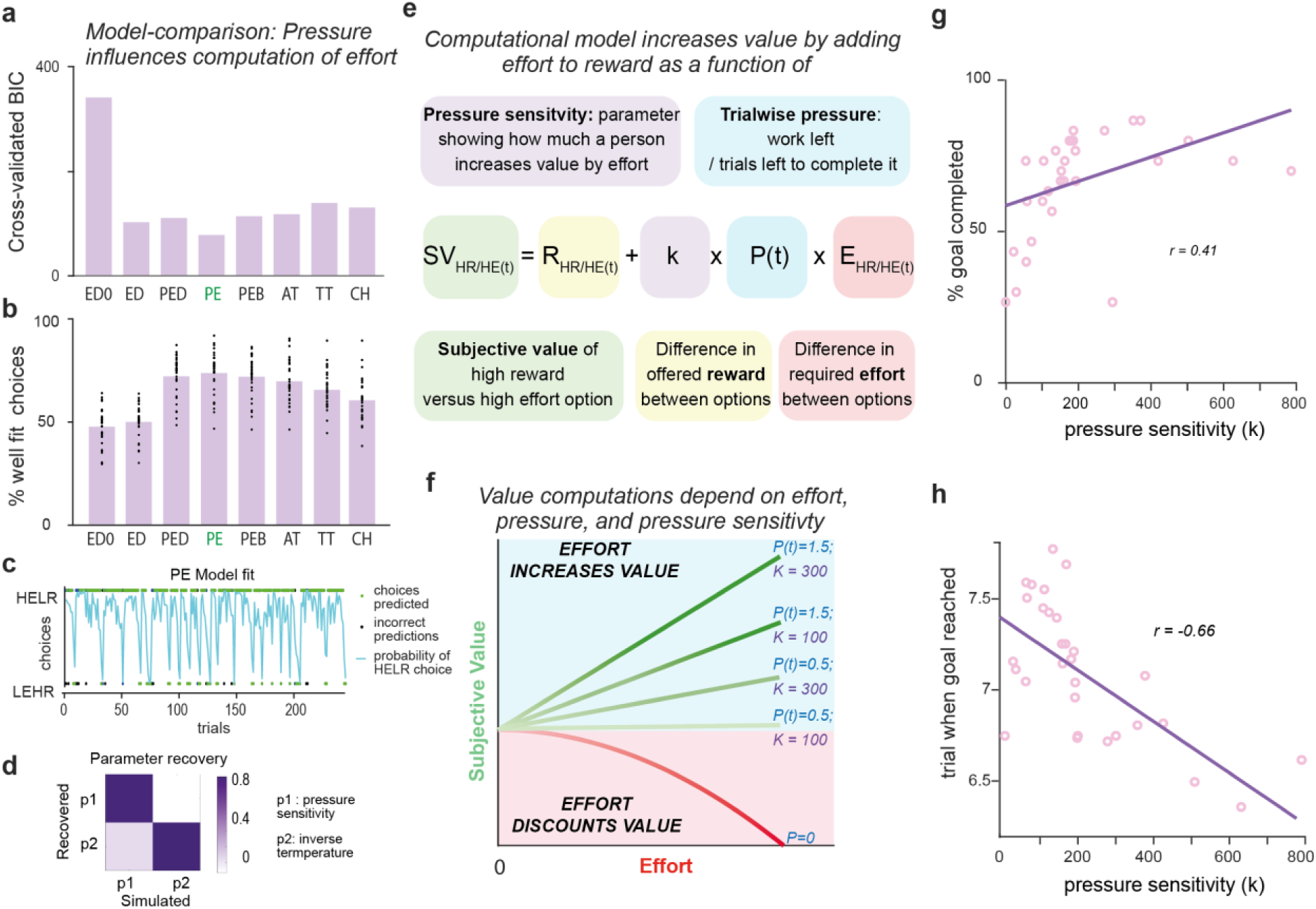
Computational modelling of deadline pressure and the valuing of effort. **a**. Cross-validated BIC values for different computational models of decision-making in the main task (lower is better model fit). The winning model is PE shown in green. **b.** average percentage of correct choices across models. The winning model predicts on average 74.1% of choices. **c.** an example of fit of the winning model for one participant, predicting 91% of the participant choices. The model computes a probability (cyan) of higher effort lower reward choice for each of the 240 trials (30 blocks of 8 trials). 92% of choices were correctly predicted (green dots). **d.** Parameter recovery of model parameters from the winning model, where (P1) is the inverse temperature (p2) is the pressure sensitivity parameter. **e**. The winning model. The value of both options depends on reward, but the amount of effort offered by an option adds to the value of choosing that option, as a function of the current level of pressure (work left to complete goal divided trials left to complete it) and participants’ sensitivity to that pressure (k). **f.** Schematic representation of the winning model. The linear relationship between value and effort is modulated by pressure. For positive values of pressure, effort increases the value of a choice, for negative values of pressure, effort discounts the value. **g, h.** The individual subject pressure parameter values fit by the winning model correlate with the percentage of goal reached by participants (f) and negatively correlated with the trial at which goals were reached on average. Each dot is a participant.

We find across all three studies that the best fitting model according to cross-validated BIC, was a model where the pressure on a given trial increases the value of effort (BIC = 249.7 for study 1, BIC = 248.0 for study 2, BIC = 332 for study 3, **Fig.3b**). This model was the most successful in predicting choices (**Fig.3b; Suppl. Fig. 5**; **Fig.3c**). Strikingly this model is able to capture effort seeking under pressure with only two parameters: deadline pressure sensitivity and stochasticity. Both parameters were successfully recovered well and independently of the choice stochasticity parameter in simulations (Methods; **Fig 3d**). According to this model, pressure acts as a scaling factor that modulates how valuable effort becomes when useful to reach a goal (**Fig.3e**). As pressure approaches zero there is a switch in the computation guiding the choices, and standard effort-discounting returns as is typically found (**Fig.3f**). Thus, crucially our model corroborates the statistical analyses above, showing that pressure drives people to value effort.

## People who were more proactive reached goals more often and earlier

Having established that pressure underlies choices, we next wanted to test whether it was associated with variability between people in preferences for when to exert effort to reach goals. One possibility is that people aim to reduce pressure before it builds, and thus choose higher effort more often in earlier trials. Alternatively, people might let pressure build before choosing higher effort closer to the deadline. To examine individual differences, we focused on studies 1 and 3, as study 2 had too small a sample to reliably examine variability between subjects. There was some evidence that participants were proactive, making more effort and choosing the higher effort option more in the first half of blocks although this didn’t replicate in study three (Overall effort study 1: *t* = 6.39, p <0.001; study 3: *t* = 1.25 *p* = 0.22; first 4 trials vs last 4 trials study 1: *t* = 4.85 *p*<0.001; study 3: *t* = –0.37, *p* = 0.71; Fig.2a; **Suppl. Fig. 2; Fig.2a**). However, strikingly, variability in choosing the higher effort option on the first trial was highly correlated with how many trials into a block it was when they completed goals on average (r = – 0.73 p<0.001 for study 1; r = – 0.42 p<0.01 for study 3). Thus, people who were more proactive and chose the higher effort option more on earlier trials, reached goals earlier.

The winning computational model included a single parameter that can account for variability in the sensitivity to pressure, scaling the extent to which pressure leads to an increased value of effort (**Fig.3e and 3f**). We found that this parameter was associated with a higher proportion of goals reached (r = 0.41 p<0.01 for study 1, fig.3g, although not reaching significance for study 3: r = 0.20 p=0.06), earlier reaching of goals (r = –0.66 p<0.001 for study 1, fig.3h, and r = –0.52 p<0.001 for study 3), with lower levels of average experienced pressure (r = –0.54 p<0.001 for study 1 and r = –0.62 p<0.001 for study 3), and a higher proportion of higher effort choices in the first trial (which have equal pressure for all participants on average across all blocks) (r = 0.76 p< 0.001 for study 1; r= –0.76 p<0.001 for study 3). Interestingly, in study 3, it was also negatively correlated with self-reported daily levels of fatigue (r = – 0.35, p < 0.05, **Suppl. Fig.5c**). This suggests that the computational mechanisms underlying deadline pressure may be linked to the willingness or ability to exert effort to reach goals before deadlines in everyday life.

These behavioural results provide a computational account in which the pressure exerted by a deadline, leads to higher effort being valued and chosen, rather than being an avoided cost. Moreover, they show that sensitivity in the willingness to proactively (**Fig. 2f/3h**) reduce pressure may lead to earlier goal attainment and may be a marker of everyday exertion of effort.

## MCC and putamen signal deadline pressure during choice

We predicted that regions of the brain previously linked to effort-based decision-making, and the tracking of task progress, would integrate information about the time left before a deadline and the amount of progress left to complete, to signal the pressure to exert effort. That is, the BOLD signal in some of these regions would fluctuate trial-by-trial with the pressure experienced by the participants when deciding how much effort to exert in order to reach the goal. To test this notion, we fitted trial-by-trial values of deadline pressure time-locked to the moment when participants were presented with the offers (**Fig.4a**). We found (**Supplementary Table 1**) a significant negative relationship between pressure and the BOLD signal in a cluster extending across the MCC, with the peak inside the posterior rostral cingulate zone (RCZp^36^, peak MNI: –1, 4, 35; Z = –5.66; p < 0.05 FWE cluster-corrected; **Fig. 4b**). Notably this MCC cluster did not contain voxels significant for chosen effort or subjective value. This cluster was uniquely associated with pressure and did not respond to the advancement towards the goal or the trials left to reach the goal (**Fig. 4**). Additionally, we found a cluster associated with the same pressure variable in the putamen (peak: 33, 0, –4; Z = –5.80, p < 0.05FWE cluster-corrected; **Fig. 4c**). Thus the putamen and MCC, which are regions previously linked to effort-based decision-making, showed variability in their responses as a function of the level of pressure on each trial.

**Figure 4.**
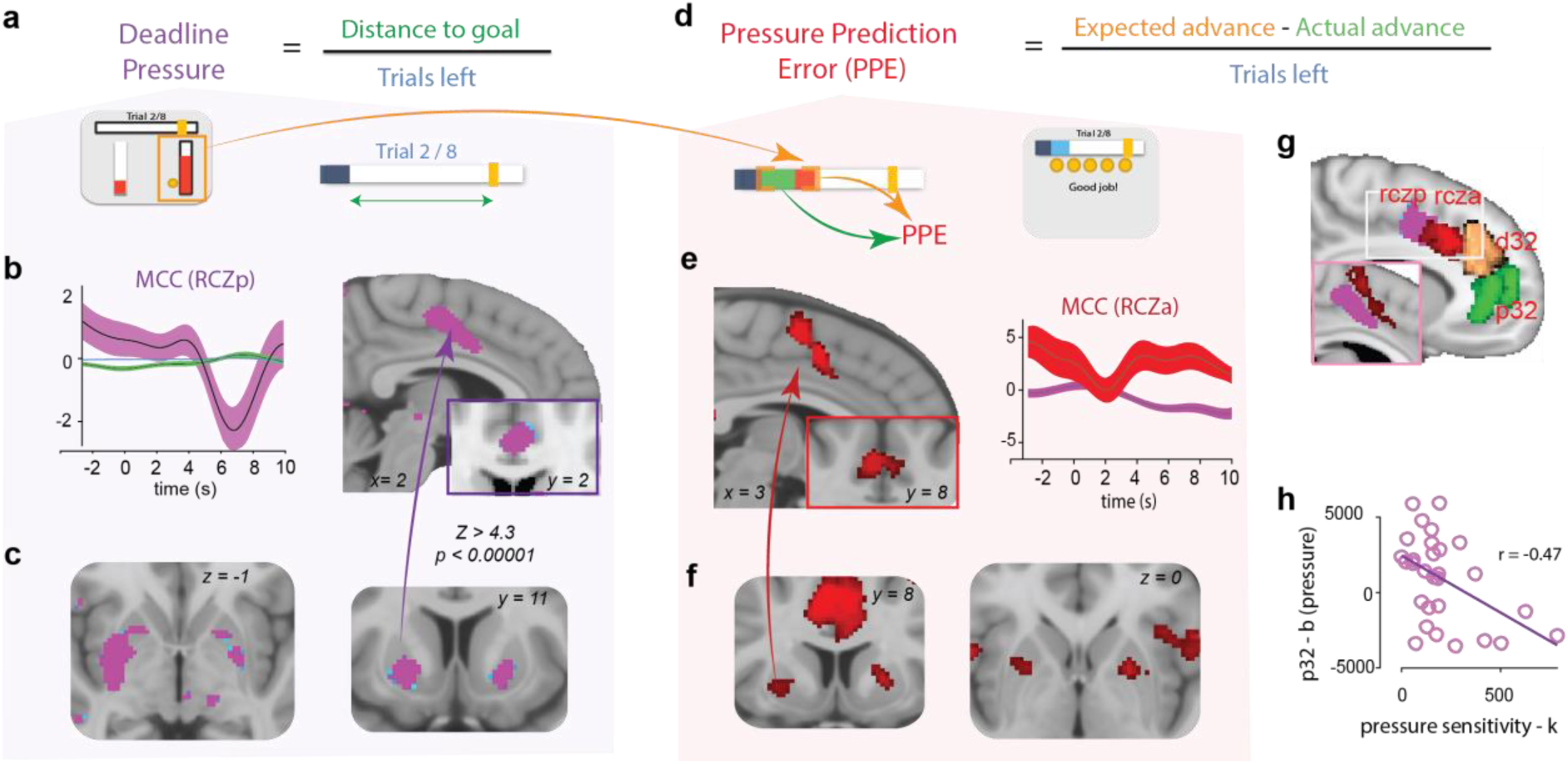
Neural responses to pressure and pressure prediction error. **a**. Deadline pressure was computed as the ratio between the distance to the goal (green) and the time left to reach it (blue), that is, the trials left before the end of the block. **b.** BOLD response signalling trial by trial pressure at the time when participant saw the offers in the MCC (purple). In the left panel, time series of the strength of pressure (purple), distance to goal (green) and time left (blue) representations in voxels from an independent anatomical ROI in RCZp. **c.** BOLD response signalling pressure in the Putamen. A PPI analysis using the posterior Putamen as a seed revealed a significant increase in connectivity at time of offer in MCC. **d.** As the mapping between effort and advancement was probabilistic, in each trial subjects experienced a discrepancy between how much they expected to advance (orange) based on their choice (orange arrow) and what they actually advanced (green). Therefore, subjects could experience a pressure prediction error (PPEs) weighting this discrepancy by the time left. **e.** BOLD response signalling trial by trial PPEs at the time when participant saw the outcome in the MCC (red). In the right panel, time series of the strength of pressure and pressure prediction error representation at time of outcomes in voxels from an independent anatomical ROI in RCZa. **f**. BOLD response signalling trial by trial PPEs in the Putamen (shown at z>3.1). A PPI analysis using the posterior Putamen as a seed revealed a significant increase in connectivity at time of feedback with MCC. **g.** Four independent anatomical masks for sections of the MCC and ACC. RCZp and RCZa broadly overlap with the peaks of the adjacent representation of pressure and pressure prediction errors. **h.** Correlation between the pressure sensitivity parameter and the average βs in ACC (anatomical region p32). Each dot is a participant.

Next, we performed a psychophysiological interaction (PPI) analysis time-locked to offer presentation with seeds in a posterior putamen mask (Suppl. Fig. 7) and in a mask of the RCZp which were anatomically defined to test where these regions might functionally connect with during choice. This identified a cluster overlapping with the RCZp cluster responding to pressure was identified from the putamen PPI seed, and vice versa a putamen cluster was present in the RCZp (**Fig. 4c and Suppl. Fig.7c**), indicating the two regions were functionally connected during decision-making. Thus, at the time of evaluating and making effort-based decisions, activity in a connected network including MCC and Putamen encoded the pressure that is experienced by participants while working to reach a goal.

## MCC and putamen signal deadline pressure prediction errors during outcomes

A crucial component of working towards goals is that sometimes, for the same amount of effort, we make more or less progress than we expected. We therefore hypothesised that this could be conceptualised as a pressure prediction error (PPE) at the time of the outcome of effort during the feedback screen. That is, we exert effort that we believe will reduce pressure to a certain degree, and if the outcome of our effort reduces pressure more or less than this, the difference can be considered as a PPE. In our task, by manipulating the mapping between effort and advancement towards the goal in different blocks (i.e. high and low pressure blocks), we created uncertainty about the amount that pressure would be reduced by an effort on each trial. Thus, in each trial, we could calculate a signed PPE and look for areas in which activity was scaled trial by trial with PPEs. We found (**Supplementary Table 1**) a cluster in the MCC in which activity covaried with the PPE (peak: 3,9,46; Z = 4.93, p < 0.05 FWE cluster-corrected; **Fig. 4e**; **Fig. 4g**). Notably while this cluster was also in the MCC, the cluster appeared to be anterior to the RCZp, lying in the anterior portion of the RCZ extending across the anterior rostral cingulate zone (RCZa^36^). In addition, we found a significant cluster in the putamen (peak: 22, 4, 2; Z = 3.76, p < 0.05FWE cluster-corrected; **Fig. 4f**). While this cluster was partially overlapping with the putamen cluster signalling pressure at the time of offer presentation, its peak did not fall within that other cluster. This suggests that there may be anatomical separation in the putamen as well as in the MCC between different sub-regions signalling pressure and PPEs. PPI analyses at the time of the feedback, also revealed clusters in the MCC from the posterior putamen seed, and vice versa, suggesting these regions are functionally connected when monitoring and updating the level of pressure to reach a goal (**Fig. 4f and Suppl. Fig.7c.)**

## Distinct portions of ACC signal effort when under deadline pressure or not

Previous research has shown that across the MCC, ACC, VMPFC, Insula and basal-ganglia effort and reward are processed when people evaluate whether to exert effort or not. Typically, they are found to show responses in opposing directions, i.e. if activity increases with reward magnitude it reduces with higher effort^10^. However, as participants showed a pattern of behaviour reflecting the positive valuation of effort, we aimed to examine if similar or distinct regions signal effort when it is being treated as a cost or when it is being processed as valuable. To do this we compared trials where participants had yet to complete the goal (pressure trials) with trials where participants had completed the goal (no pressure trials). Behaviourally participants switched back to avoiding the higher effort on no pressure trials, as they would in an effort discounting task, as there was no need to make more progress. We then looked for activity that covaried with the chosen effort on those two trial types.

This revealed two distinct clusters in the anterior cingulate cortex (ACC). On pressure trials where effort adds value, BOLD responses varied with chosen effort in the rostral ACC, in the ventral tip of area 32 (p32, peak:-1,30,-3; z = –4.11, p < 0.05FWE cluster-corrected; **Fig. 5a**). In contrast on no pressure trials BOLD responses in a more dorsal ACC cluster responded to chosen effort, in a region more commonly found in studies examining effort-discounting processes (d32 extending to RCZa, peak: 6,35,22, Z=-4.10, p < 0.05FWE cluster-corrected; **Fig.5a**). These results corroborate the idea that chosen effort processing differs when there is no pressure, compared to when people are under the pressure of a deadline. Once more a cluster in the right posterior putamen signalled effort both in trials with or without pressure (peak without pressure: 29, –4, –2; Z = 3.97; peak with pressure: 29, –3, –5; Z = 4.69, p < 0.05 FWE cluster-corrected; **Fig. 5b-c**). However, it appeared that the putamen processes effort on both types of trial in the same manner (**Fig. 5b-c**), suggesting that processing the effort of the action was the same in the putamen regardless of whether it was being treated as a cost or valued.

**Figure 5.**
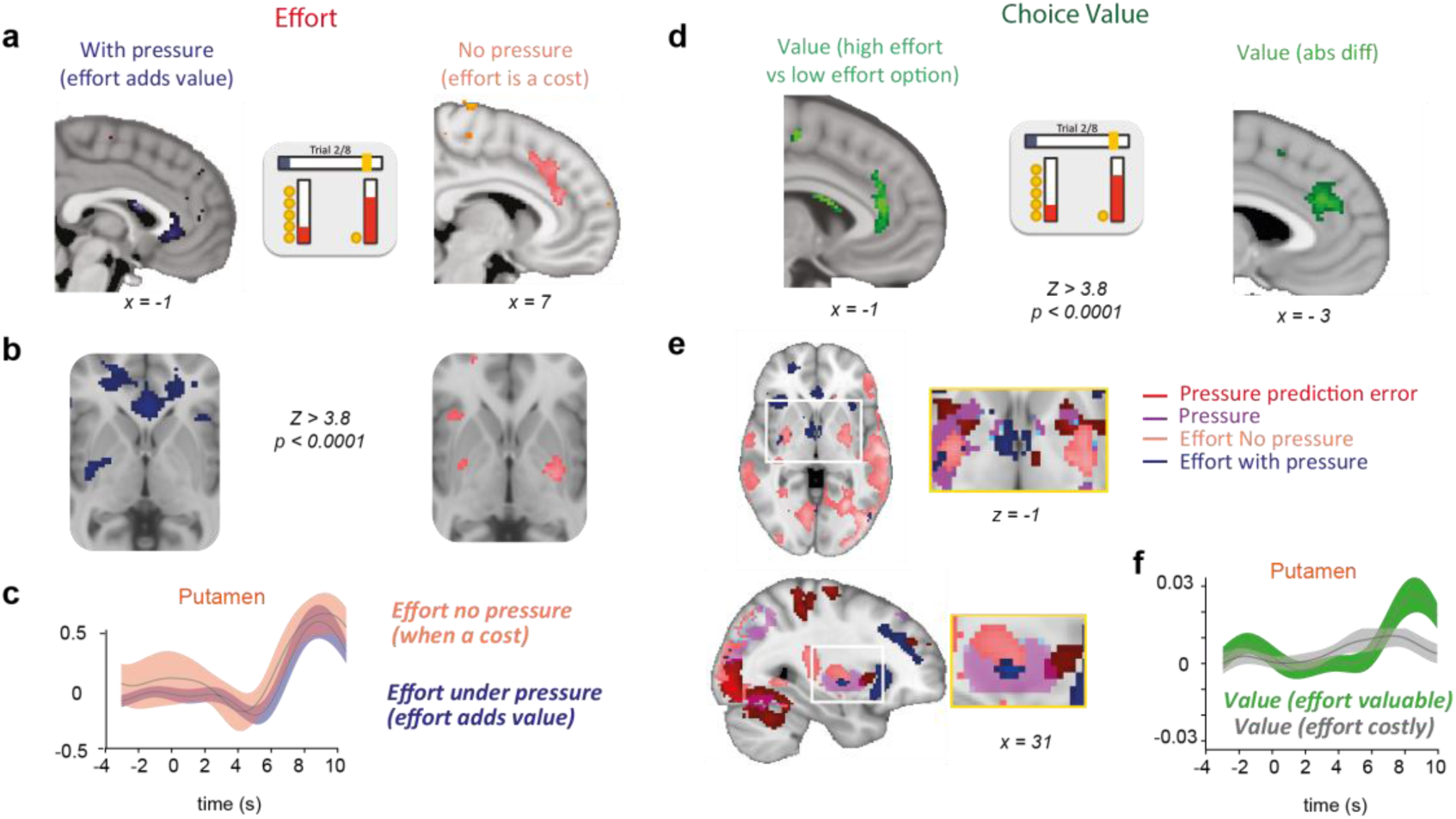
Neural representation of effort and value. **a**. Representation of effort at time of choice in the ACC in trials where subjects experienced pressure (blue) and in trials when they didn’t (pink). **b** Representation of effort in the putamen. **c**, Time series of the strength of effort representation in voxels from an anatomical ROI in the posterior putamen. **d**. Representation of the difference in value between the offers at time of choice in the ACC. Signed difference in value between higher effort and lower effort option (left panel) and absolute difference in value between the options (right panel). **e.** Representation of effort, pressure and pressure prediction error in the putamen. **f**, Time series of the strength of value representation in voxels from an anatomical ROI in the posterior putamen.

## A different ACC subregion signals subjective value

Next, we wanted to test if, and where, subjective value from our pressure model (in which effort increases value) was processed. To test this, we took the trial-by-trial estimate of the difference in value between the offers according to the model, and examined activity at the time when participants were presented with offers. We found a significant relationship between the BOLD signal in a dorsal ACC cluster extending across the dorsal portion of area 32 (d32 peak: –3, 36, 15; Z = –4.11, p < 0.05FWE cluster-corrected; **Fig. 5d left panel**). In a separate analysis we used the absolute difference in value between the two options and found a significant cluster in area 32 (d32 peak: –2, 37, 26; Z = –5.15, p < 0.05FWE cluster-corrected; **Fig. 5d right panel**). Both these closely related representations disappeared in a control analysis where subjective value was computed from a standard effort discounting model which is not informed by pressure (ED0 – see equation 2 in methods), suggesting these regions were processing value only when effort was positively valued.

Next, we extracted ROIs in four anatomical regions^36^ of the cingulate sulcus and tested whether the strength of their representation of pressure and pressure prediction errors were correlated with the pressure sensitivity parameter independently estimated for each subject by the model. That is, do any of these regions show a relationship with how sensitive to pressure a participant is. We found a negative relationship between the encoding of pressure in the p32 region of the cingulate cortex and the pressure sensitivity parameter (r = – 0.47 p < 0.05; **Fig. 4h**). This cluster overlapped with that which signalled chosen effort on pressure trials where effort was having a positive value. This suggests that this is involved in changing the value of effort to positive as a function of how sensitive a person is to deadline pressure.

We also, once again, found a cluster in the putamen, this time, both for the absolute difference in value incorporating pressure or in the model not including pressure (peak for value difference with pressure: 31, –2, 4; Z = 3.35; peak without pressure: 15, 9, –10; Z = 3.11; cluster-forming threshold *p* < 0.001; **Fig. 5f**). It was also activated by the chosen reward but slightly earlier in time (Suppl. Fig. 7). Indeed, adjacent subregions of Putamen encoded reward, effort both under – and without – pressure, pressure, pressure prediction errors and subjective value, suggesting that Putamen acts as an integration hub of these different computations allowing a switch in the value of effort (**Fig.5e and Supplementary** Fig.7).

## Discussion

Using computational modelling and ultra-high-field fMRI we revealed neural and computational mechanisms underlying decisions to exert effort to reach goals under the pressure of deadlines. We replicated in three experiments the finding that people seek higher effort behaviours, that are associated with lower immediate rewards, because the pressure caused by the deadline associated to a goal (how much progress towards the goal is left to complete divided by the time left to complete it) switches effort to be valued rather than being treated as a cost. Activity in portions of the MCC signalled this deadline pressure with a neighbouring region of MCC signalling pressure prediction errors when the outcomes of an effort reveal pressure had reduced more or less than expected. We found representations of the subjective value of rewards with the added value of effort, effort when it is treated as a cost, or as a benefit in three separate ACC sub-regions. The putamen was also found to signal all these five decision variables. These results combined highlight how the brain processes the pressure caused by deadlines, how this shifts the valence of effort within decision-making, and switches processing in multiple circuits in the brain that keep people on track to meet goals.

Deadlines are a ubiquitous feature of modern societies and foraging environments^1,2,4,6,24^ and a powerful motivator of effortful behaviour^37^. However, existing research had not provided an account of the neurocomputational mechanisms underlying this phenomenon. We offer evidence for a normative account for how and why deadlines increase effortful exertion, that may explain multiple existing findings in motivation research.

Firstly, we suggest effort can be valuable when it makes us progress towards goals. While participants were instructed that greater effort made more progress towards goals, it may also be that people can learn this (e.g. learning that effortful training improves performance^38,39^) and generalize it from one context to another. This may explain why, despite effort being typically considered aversive, in circumstances where people have set a goal, they seek it (e.g. cycling up a mountain). Notably, participants did not need to value effort to achieve goals in our task. It was necessary to choose the higher effort option to succeed, but goals could be reached through other strategies. For instance, choosing high effort until the goal was reached regardless of the amount of offered effort and reward, or choosing the higher effort option only when it was at lower effort levels. These would be successful strategies but were not apparent in behaviour. This concords with existing research showing that people can value some effortful actions, and show increased liking of objects they have exerted effort into making (the Ikea effect), and learned industriousness effects^22,26,40^. However, we demonstrate a new circumstance, and a normative account for how goals with deadlines can lead to positive valuations of effort.

Secondly, we propose the switch in valence of effort occurs due to the pressure of having to complete a significant amount of effort to reach a goal before a deadline. This offers a mechanistic understanding for why deadlines/finish lines can increase motivation and performance^41^. This goal gradient effect^41^, had been argued to be due to opportunity cost-based mechanisms, i.e. finish lines make effort seem more worthwhile^24,42^ or increase the marginal value of progress^43^. However, this line of work could not determine whether such effects were because people’s choices of whether to exert effort were changing due to a reduction in the perceived cost, or because effort was being valued, nor offered a computational account for why this change in value might happen. Our notion of deadline pressure can putatively account for these effects.

Thirdly, while we show effort can be positively valued in choice, it does not necessarily mean that it was hedonically pleasurable^22,44^, nor that there was not a cost to that effort for the motor system^45^. Indeed, we found that the putamen signalled chosen effort on all trials regardless of whether it was trials where effort was valued or costly within decision-making. Thus, it is plausible that the brain can simultaneously represent a different valence for the same act in different brain systems, potentially explaining why it may appear paradoxically valued at some points, but not at others.

Lastly, our results are consistent with the idea that avoiding uncertainty and risk of missing goals could be an overarching motivation. In our task maximising reward required succeeding at goals. Thus, it would be risky not to choose the higher effort option as avoiding effort would mean not progressing and subsequently failing. Thus, deadline pressure could in theory be considered as a calculation of the risk of not maximising long-run reward. This would suggest an overarching driver of the choice of higher effort may be the uncertainty over whether important long-term goals will be achieved, which has previously been shown to increase risk-taking^34^. Deadline pressure, and its effects on effort, may therefore reflect a crucial component for maximising reward in foraging animals^46^.

Interestingly some people were highly proactive and sensitive to deadline pressure, choosing the high effort option frequently, early on during goals. This variability was captured by the pressure sensitivity parameter within our model which in turn was associated with daily levels of fatigue. Previous work has shown that higher levels of fatigue are associated with a reduced willingness to exert effort for reward, putatively due to an increased cost of effort^47,48^. We extend this, suggesting that feelings of fatigue prevent people from being able to proactively respond to deadline pressure and treat effort as valuable. Moreover, this suggests that our computational model may be fruitful for understanding variability in motivation^49,50^, and disorders associated with reduced fatigue^16,17,51^.

The finding of high levels of proactivity overall could be interpreted as being inconsistent with people procrastinating and not focusing on goal completion. Procrastination often occurs in the presence of alternative goals which distract from a main task because they can offer more gratification^52^ or are perceived as more urgent^53^. Without an alternative goal, participants thus had little else to procrastinate on in our study. However, it is also important to note that participants did fail on multiple blocks. Previous research has shown that procrastination may occur when people are tempted by an immediate reward^54^. As people were presented with a low effort (low progress) high reward option, it is plausible that this caused a degree of procrastination leading to failures to complete goals. Future work should try and disentangle how this lure of alternatives and deadline pressure interact to lead to failures to reach goals due to procrastinating.

This work provides the first characterisation for how sub-regions of ACC, MCC and basal-ganglia might guide our motivation to exert effort to succeed in reaching goals before deadlines. The MCC has been found to signal the costs of effort during effort-based choice^10,21,32^, unexpected changes in required effort^55^, for motivating people to overcome effort^16,56^, to persist with effortful behaviours^57,58^, and also signalling fluctuating levels of fatigue during effort-based choices^23,51^. Other studies have also suggested the MCC may be important for signalling wider contextual information that influences other types of choice and cognitive control processes^33,59–62^. This includes the pressure to make risky choices to reach goals^34^ and the tracking of progress through sequential tasks^9,63^. The anatomical connectivity of this region places it as an area that receives input from sensorimotor, interoceptive and cognitive processing areas^36,64,65^. Moreover, it contains neurons that process unexpected outcomes, monitor errors in task performance, and lead to post-error slowing in cognitive control tasks^66–70^. With our results suggesting it signals deadline pressure, this paints a complex picture for what function this processing may serve.

Our model of deadline pressure can potentially encompass and integrate these previous findings regarding MCC function. Our results suggest the MCC integrates contextual information about how aligned current progress is with respect to one’s goals, that provides a function (deadline pressure) at the time of choosing that indicates the need to exert effort to reach a goal, and another (pressure prediction error) to monitor and update this over time. These computations, in addition to those we identified within the ACC and putamen, allow individuals to flexibly update goal-directed motivation and behaviour to ensure our efforts allow us to reach our goals over time. This therefore provides a framework for a more rich, complex approach to motivated behaviour, that may be applicable to multiple other forms of cognitive control and performance monitoring processes linked to MCC function, where people have longer-term goals.

Overall, our results in the putamen suggest it is involved in integrating multiple features that guide decision-making (subjective value, effort cost, effort benefit, pressure and pressure prediction errors) through communication with the anatomically connected MCC and ACC^71,72^. This is supported by previous evidence showing that the putamen is engaged when making effort-based decisions, with variability in its responses linked to effort aversion^28,73^. Notably, both when treated as a cost or valued, effort and the subjective value according to the model, were found in the caudal putamen. Moreover, the putamen signalled effort on all trials in the same manner, regardless of whether people will still aim to achieve a goal or it had already been completed. This is in line with existing evidence that suggests more posterior portions of the putamen guide automated motor responses and force exertion^30,73^. The posterior/caudal putamen may therefore be more directly involved in processing the cost of exerting effort regardless of whether it was positively or negatively valued in choice.

In contrast, we found that pressure and pressure prediction error signals appeared to be localised more anterior/rostral within the putamen. This expands existing ideas of the contribution of the putamen to behaviour, suggesting it may integrate information about the context that influences the processing of effort. With dopamine neurons in the substantia nigra projecting widely across the putamen^73–75^, this may offer an explanation for some motivational impairments occurring in disorders linked to basal-ganglia, such as Parkinsons Disease (PD)^16,76,77^. While loss of dopamine function in PD has been linked to many aspects of motivation, this includes an inability to perform tasks with multiple components^78^. Initially this was believed to be due to deficits in planning. However, as has been noted, the ability to plan is intact in PD, but execution appears to be disrupted^79^. Our results suggest that the putamen plays a crucial role in integrating information about goals and deadlines into signals that motivate individual behaviours towards longer-term goals. The signals we identified were in a task where explicit planning was not possible, instead one had to motivate different amounts of effort based on task progress. A deficit in these mechanisms could lead to the inability to motivate the exertion of effort into specific acts during a sequence of behaviours. Future research should therefore investigate how neuro-modulatory, potentially dopaminergic, and basal-ganglia related deficits might impact on how effort-based decisions are made when progressing towards a goal before a deadline.

Previously there has been debate about whether the medial frontal cortex is engaged in processing the subjective value of rewards, the difficulty of decisions, foraging value, and effort costs^10,60,62,80,81^. This has included some specific questioning about whether value during effort-based decisions is signalled in dorsal ACC^10,11,31^. Notably, we found a spectrum of computations being performed across multiple sub-regions of the MCC and ACC with the effort being chosen signalled in different sub-regions of the ACC when it was being processed as a cost or when it was being valued. Another distinct sub-region signalled subjective value according to our computational model. This would suggest that there are different mechanisms that might guide effort-based choices when aiming to reach a goal, compared to when there is no goal as is the case in past experiments, and argues against the idea that our results could be explained by one alternative factor such as choice difficulty. Our results therefore suggest there is not a single system for guiding effort-based decisions, but instead multiple systems depending on the context in which those choices are made, with this all integrated within the putamen.

Deadlines have a major impact on motivation and are ubiquitous features of many aspects of our daily lives. Using computational modelling and ultra-high-field fMRI we show that when under the pressure to reach a goal before a deadline, people switch to treating effort as valuable, rather than costly, during decision-making. People who are proactive and work hard early on are more successful in completing goals, which is more challenging for people who suffer from high levels of fatigue. A spectrum of computations across medial frontal cortex, and functionally connected putamen regions, signal deadline pressure, update this through a prediction error, signal the subjective value of effort and switch from signalling effort from costly to valued. Thus, we provide a computational framework for how fronto-basal-ganglia circuits help people keep ‘on track’ to exert effort and reach their goals before deadlines.

## Methods

Three studies were undertaken to examine how people make decisions about whether to exert effort in order to reach a goal before a deadline. Study 1 was completed while undergoing 7T fMRI aiming to examine the neural correlates, study 2 was designed to replicate the main behavioural findings, and study 3 was conducted with an online version of the task aiming to examine variability between participants in behaviour.

## Participants

We recruited 95 healthy young participants across three studies. In study 1 (N = 30, 16 female, 14 male, age range 18-40) 3 participants performed a lower number of blocks (21, 19 and 15 instead of 30) due to technical problems with the scanner. The sample size for study 1 (N= 30) was chosen based on previous effort-based studies reporting strong effect sizes obtained with 3T fMRI scanners at N = 35 ^10,27^ and considering the up two times higher signal to noise of 7T scanners^82^. In study 2, (N = 15 participants, 8 female, 7 male, age range 18-40) participants performed the task while sitting in front of a computer in an isolated room. The sample size was based on a power analysis of the effect size of pressure that a smaller sample was sufficient to replicate. For both of these studies the recruitment was from the student population of the University of Oxford, and the wider community of the city of Oxford. Thus, in study 3, (N = 40, 11 female 29 male, age range 20-45) participants were recruited from online platform Prolific.com and performed a version of the task online on Gorilla.sc, an online experiment builder. This ensured a wider sample of UK participants. The sample size for study 3 was chosen based on related previous studies assessing the relationship between fatigue scores with effort-based choices^47^ for N = 40. All studies were approved by The University of Oxford Central Research Committee. Informed consent was obtained from all participants. Four participants (2 in study 2, 2 in study 3) were excluded due to a failure to follow task instructions or for low performance in the task (the exclusion criterion was met if the participant didn’t succeed in at least 10 blocks out of 30). For the first two studies, participants were compensated a flat rate of £12.5/hour for their time and were also told that they would receive a bonus of up to £5 based on their responses in the task. For the third study, participants were compensated a flat rate of £9.76/hour for taking part in the experiment and were also told they would receive a bonus of up to £5.

## Procedure

Across all studies, the same basic design was used to examine how the willingness to exert effort to obtain rewards changes over trials under the pressure to reach a goal by a deadline. All studies comprised an initial *calibration* phase to establish their maximum voluntary contraction (MVC – study 1 and 2) on a handheld dynamometer, or maximum number of mouse button clicks (MVNC – study 3) followed by a *training phase*, in which participants were familiarised with each level of effort that they would be required to perform during the rest of the experiment. This was followed by an *effort-discounting task* to assess their effort sensitivity in a more standard paradigm where there was no goal, and the *deadline task,* where they had to complete goals.

*Calibration –* In the first two studies, we measured their maximum voluntary contraction (MVC) to normalise force levels across participants and avoid variability due to differences in strength. Participants were asked to squeeze the dynamometer as hard as they could for three times, while receiving verbal encouragement. During each attempt, a vertical bar presented on the screen provided live feedback of the force level generated. In the second/third attempts, a benchmark representing 105/110% of the previous best attempt (displayed with a “target” yellow line on the screen) was used to encourage the participants to improve on their score. Participants did not know that this yellow line indicated a higher force level than in their previous attempts. The maximum level of force generated during the three attempts was used as that participant’s MVC for the rest of the experiment. In the third study, a similar procedure was performed, asking participants to click as many times as they could on three consecutive occasions. The maximum number of clicks was used as that participant’s maximum voluntary number of clicks (MVNC) for the rest of the experiment.

The levels of force required throughout the experiment were then computed as percentages of each participant’s MVC/MVNC. We defined 4 effort levels corresponding to 20%, 35%, 50%, 70% of MVC for study 1 and 2 and for effort levels corresponding to 50%, 65%, 80% and 95% of MVNC for study 3. The difference in effort levels between study 1 and 2 compared to study 3 reflect the inherent difference in effort between the two tasks employed, whereby clicking requires proportionally less effort than squeezing a handgrip.

*Training –* In the first part of training, participants were familiarised with how much effort was required at each level. Participants were presented with the 4 effort levels in an ascending order. In the first two studies, for each effort level, a yellow line was shown representing the required level of effort while a fluctuating vertical red bar represented the instantaneous effort exerted by the participant on the hand-held dynamometer. Participants had 3 seconds to reach the required effort threshold by squeezing the dynamometer. The response was considered correct if the effort exerted was above the required level for at least 1, non-continuous second, out of 3 seconds. The same procedure was adopted for the exertion of all levels of effort throughout the rest of the experiment. For study 3, a round grey button was shown for 3 s and had to be clicked over until the required number of clicks. The same procedure was adopted for the exertion of all levels of effort throughout the rest of the experiment.

In the first part of the training, participants were asked to reach the required level of effort to complete each trial in order to gain one credit. Thus, participants were familiar with how much effort would be required for each level prior to making decisions in the main task.

In the second part of training, participants were introduced to the kind of choices they would experience during the main task. Participants were asked to complete a block of 8 trials, and in each trial choose between two options with different levels of reward and effort, a highly rewarding option that required little effort and a very effortful option which was not very rewarding. In the first two studies, effort levels were represented by a vertical red bar, while reward levels by a pile of coins next to the effort bar. In the third study, effort levels were represented through a pie chart with increasing number of slices (from 1 to 4), while reward levels were represented by a horizontal pile of coins under the effort pie chart. Participants indicated their choices by pressing one of two keys on the keyboard to select either of the two options, presented on the corresponding left or right of the screen. Participants were also shown a progress bar with a goal they would need to reach by exerting enough effort over the course of 8 trials. In this part of the training participants were familiarised with how to advance towards the goal in the main task by the exertion of effort.

## No-goal (effort-discounting) task

In order to test whether the participants we tested showed a typical aversion to effort, participants performed 30 trials of an effort-based task which did not include a goal and was therefore performed without any pressure. Here, each trial was independent of each other and involved a choice between two options associated with different level of reward and effort. Taking this approach allowed us to measure the degree to which people devalue rewards by effort in a task where they were not incentivised by the need to reach a goal. In the pre-task, the work options were sampled from four reward levels (1, 4, 7, 10 credits) and four effort levels; for the task in the first two studies this corresponded to force (20, 35, 50, 70% MVC). In the third study this corresponded to number of clicks (50, 65, 80, 95% MVNC). Effort and reward levels were chosen based on pilot and previous experiments ^10,27^. This task was always performed before the main task below, to ensure it was unlikely to be affected by fatigue or having performed the main task, that might decrease the willingness to exert effort. While in the main task all choices were followed by the requirement to exert force, in the pre-task only 20% of choices required effort exertion after choice (50% for the third study) in order to reduce the experimental time and reduce effects of fatigue. Participants indicated their choices by pressing one of two keys on the keyboard to select either of the two options, presented on the corresponding left or right of the screen. The trials requiring effort were pseudo-randomly selected.

## Deadline pressure task

In the main task of the experiment participants were tasked with trying to reach a series of goals, each of which needed to be completed within a block of 8 trials. To progress towards the goal, subjects had to exert effort, where more effort led to a greater amount of progress. If they reached the goal, their bonus payment for participating would increase as a function of the number of coins collected during the task. If they did not reach the goal by the end of the 8^th^ trial, they did not obtain any of the credits they had collected. The amount of effort they could exert on each trial depended on choices between two options. On each trial they would be required to choose between an option that would make more progress (higher in effort) but would obtain less reward (coins), or an option that was lower in effort that would make less progress but higher in reward. As such, participants would fail at this task if they avoided the higher effort, lower reward option.

Each trial began with the presentation of two choice options. Each option comprised an effort level, represented by the height of red bar fill in the first two studies and by the number of slices in a pie chart in the third study, and a reward magnitude, represented by the number of coins next to the effort bar or under the pie chart. Participants were always presented with a “high(er) reward – low(er) effort” choice option on the left of the screen, with one of the highest reward values (4, 7 or 10 coins) and of the two lowest effort levels (20% or 35% of the MVC or 50% and 65% of the MNVC). On the right side of the screen there was a “high(er) effort – low(er) reward” (HE-LR) option, i.e. the lowest reward values 1 and 4 and the two highest effort levels (50% and 70% MVC or 80% and 95% MVNC). Participants had to make a choice between these options within 4 seconds. Failing to do so led to a “missed trial”, in which no rewards were collected, but also no progress was made, and the experiment moved onto the next trial of the block. Responses were recorded by pressing the left or right arrow buttons on a button box (study 1 in the scanner) or on a keyboard (study 2 – behavioural only and study 3 – online). After making a choice the participants had to exert the effort level chosen. Participants had three seconds to reach the required level of effort. For a trial to be considered successful, participants must maintain their force above the target line for at least one second out of three in the first two studies. A failure to do so resulted in zero progress being made, and no rewards being collected. After a jitter (1-3 seconds) feedback was presented, indicating success or failure of the trial, for 1.5 seconds. In case of successful completion, participants were shown the amount of rewards associated with their choice and the progress towards the goal.

Goals were reached only when subjects advanced enough to reach the target line on the progress bar. Advancement depended on the amount of effort exerted following the choices made by the participant: the more effort was chosen and successfully completed in a specific trial, the more advancement was made. In the first two studies, the precise mapping between the effort exerted and advancement towards the goal was probabilistic. The total amount of effort required to reach a goal was calculated as

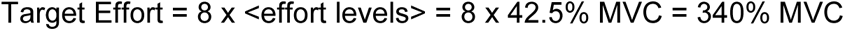

So, to progress at a pace sufficient to reach the goal in time, participants had to perform on average 42.5% MVC in each trial. However, in each trial the mapping of effort into advancement was either enhanced or reduced by 25%. In half of the blocks there was a 75% chance of enhancement and 25% chance of reduction. These were therefore easier or low-pressure blocks, where subjects would progress faster towards the goal. In the other half of blocks, the probabilities were reversed, with a 75% chance of reduction and 25% of enhancement of the mapping of effort into advancement. Therefore, these were harder or high-pressure blocks. Participants were unaware whether each block was high or low pressure and were not signalled in advance whether the mapping of effort into advancement in a trial was going to be enhanced or reduced. This manipulation made sure that participants could not plan with certainty their progress towards the goal and thus might have to choose the higher effort more than could be expected in some blocks.

Each trial began with a presentation (Fig. 1) of the offer screen in which there was a progress bar (with blue fill indicating the amount of the goal completed and a yellow line indicating the goal), the number of trials through the block (t/8), and the two effort options indicated by vertical bars with red fill indicating the effort required and yellow coins indicating the reward magnitude. After 4s the chosen offer was highlighted with a green outline, followed by the force exertion period (where real-time feedback was presented with the red filling up the vertical bar as a function of grip force). There was then a variable jitter (1-3s) followed by the outcome screen. On the outcome screen participants were shown the amount of new progress they had made towards the goal, and the number of coins obtained. This was followed by another variable jitter (1-3s). Note that these jitters were replaced with 1s fixed intervals in study 2 and 3.

At the end of each block, participants indicated how tired they felt by moving a cursor along a horizontal bar ranging from 0 (not tired at all) to 100 (extremely tired). They had a fixed five-second period to register their response using the left and right arrow keys, with the initial position of the cursor positioned on their previous rating. In study 3, before subject completed the task online they also completed a self-report 20-time questionnaire designed to measure daily levels of fatigue (Multidimensional fatigue inventory)^83^.

## Apparatus

Physical exertion was measured using a hand-held dynamometer (SS25LA, BIOPAC Systems, USA) that participants squeezed using their dominant hand. The task was programmed and presented using MATLAB 2012 (MathWorks, USA) and Psychtoolbox (http://psychtoolbox.org). All choices and responses were made on the keyboard using the non-dominant hand.

## Statistical analysis

First, we ran several logistic regression models to identify which experimental factors were most important in predicting choices in the main task. To build the best fitting model of participants’ choice data, we tested eight different models. These models included the information/elements participants viewed on the screen during task performance, such as: advancement towards the goal (Adv); number of trials left (TL); reward value of the higher reward lower effort choice option (HR); reward value of lower reward higher effort choice option (LR); effort level of higher reward lower effort choice option (LE); effort level of the lower reward higher effort choice option (HE) the pressure exerted by the goal which we defined as

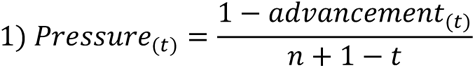

Where *n* is the number of trials, *t* is the current trial and *advancement*(t) is the proportion of total progress towards the goal that was made until trial *t*. Trials where subjects have reached the goals and *advancement* > 1 are considered “no pressure” trials in subsequent analysis.

Finally, we tested models which included the interaction between the options presented on the screen and the interaction between Pressure and the individual options on the screen. Models were compared using BIC. The best model turned out to be the one including only the values of the rewards and efforts of the two choices and the value of pressure.

## Computational modelling

To study and compare fluctuations in the motivation to exert effort under the pressure to reach a goal by a deadline, we developed several computational models considering different ways in which the presence of the goal might impact on the decision. In addition, we used computational models to examine decision-making in the task with the absence of a goal.

## Modelling choices in the absence of a goal

We first fit choices in the pre-task to estimate the choice model parameters in the absence of a goal. We used an effort discounting model which has been extensively used to accurately characterise how people devalue rewards by the effort associated with them^27,35,84^. The model assumes that the subjective value of a choice is proportional to the reward on offer, discounted by the effort associated with it. That is, effort is assumed to be a cost in such models. We used a parabolic discounting function known to describe physical effort discounting in a large number of previous studies ^27,84^:

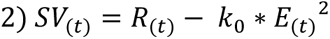

Here we estimate the subjective value (*SV*) of a choice as the value of the reward (*R*) offered on a trial discounted by the associated effort (*E*). This trade-off is dictated by a free parameter (*k*_0_) estimated for each participant. As *k*_0_ increases, it leads to lower subjective values. To fit choices between two options associated to different level of efforts we used a *softmax* transformation:

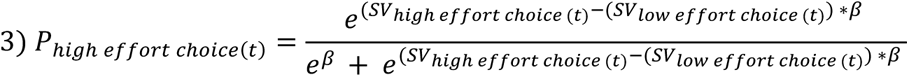

This estimates the probability (*P*_*high effort choice*_) of the participants choosing to accept the offer with the highest effort to obtain the reward. β is a free parameter that estimates the degree of stochasticity present in participants’ choices. The probability of the participant choice under the model is therefore defined as

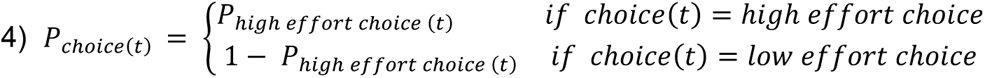

Fitting this model to the choices in the no goal task we could estimate the two free parameters (*k*_0_, β) for each participant using the *fminsearch* function in MATLAB. Each model was fit 20 times using different parameter starting values to ensure that the optimisation function had not settled on a local minimum. The parameters of this model allowed us to estimate the degree to which participants devalued rewards by effort in the absence of a goal. *k*_0_ and β were then used as fixed values when fitting some of the models to the main task data.

## Modelling choices in the presence of a goal

To fit choices in the main task we formulated different hypothesis about the effect of the goal with a deadline on decision making. As a benchmark, we considered the effort discounting model of equation 2 and fitted to the main task data (labelled “ED” for effort discounting model) and an effort discounting model with the parameters fit in the no goal task (labelled “ED0”) Both models did not compute information about pressure and advancement. We then considered different ways in which pressure and advancement might impact the evaluation of the offers. First, we considered an extension of the effort discounting model which included a term dependant on pressure to compute the value of an offer as in

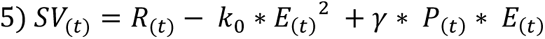

This model (labelled “PED” – pressure model with effort discounting) therefore was identical to the standard effort discounting model for P = 0 and assumed two different terms, with opposite signs, both dependant on effort. For this model we used the k0 and β fit on the choices made in the absence of a goal. Notably this model assumed that effort discounting was occurring when pressure was above zero, but this would be counteracted by the increase in subjective value by effort and pressure increasing the value. Note that we assumed that effort when adding value would not be squared, as effort was linearly associated with progress towards goals.

We then considered the possibility that in the presence of pressure to exert effort, the effort discounting term would simply disappear in the computation of value as in

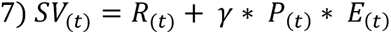

This model (labelled “PE” – pressure model without effort discounting) therefore assumed a different computation due to the presence of effort. For this model we refit the choice stochasticity parameter β.

In study 1 and 2 pressure was different on average depending on the block type. To account for this, we examined the possibility that participants would be sensitive to pressure in different ways in high vs low pressure blocks using two different pressure sensitivity parameters γ_1_ and γ_2_ depending on the block type in eq.7. This model (labelled “PEB” pressure model changing with blocks) used the choice stochasticity estimated in the no goal task.

We also considered the possibility that participants would not combine advancement and time left into a pressure value and developed a model with two separate terms as in

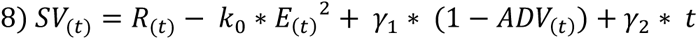

where *ADV*(t) was the advancement towards the goal at trial *t*. This model (labelled “AT”) used the effort discounting and choice stochasticity parameters estimated in the no goal trials.

We then considered the possibility that participants would only respond to time and incorporate both the past and the future time in their evaluation as in

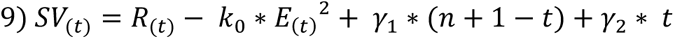

Thus affording for the possibility that participants would be proactive in looking ahead to the trials left or reactive to the trials passed. This model (labelled “TT”) used the effort discounting and choice stochasticity parameters estimated in the no goal trials.

Finally we considered the possibility that participants would ignore all the rewards and offers presented and work out their choices based on a heuristics. Up until a certain trial *q* they would choose the higher effort lower reward option 90% of the time. After trial *q* they would choose the high reward low effort option 90% of the time. *q* was therefore the only parameter fit for each participant. This model (labelled “CH” choice heuristic) had only one parameter *q*.

To compare their performance, all models were fit based on 66% of the data (training set), and the performance of each model was tested on the remaining 33% (test set). For each model, the average Bayesian Information Criterion (BIC) and the percentage of correct choices (PCC) were then computed across the three test sets (3-fold cross-validation). The BIC was computed as in

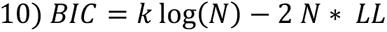

Where k was the model specific number of parameters, N was the number of choices fit and LL was the maximised Log Likelihood of the model, computed as in

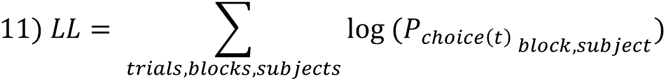

Where *P*_*choice(t)*_ was computed through equations 3 and 4 for each model.

## Model parameter recovery

We assessed the degree to which we could reliably estimate model parameters given our fitting procedure. More specifically, we generated six simulated behavioural data sets (i.e., choices for 15 high pressure and 15 low pressure blocks of 8 trials) using the parameters selected pseudo-randomly on the range obtained in the original fit. For each simulated behavioural data set we ran the winning model PE this time trying to fit the generated data and identify the set of model parameters that maximized the log-likelihood in the same way we did for original behavioural data. To assess the recoverability of our parameters we repeated this procedure 10 times for each simulated data set. The recoverability of the parameters is shown in Fig. 3d where we computed the average correlation between the parameters used to simulate the data and the recovered parameters.

## MRI data collection

We acquired the fMRI data using a 7T Siemens Magnetom MRI scanner (Siemens, Germany). Specifically, we collected functional Echo-Planar Imaging (EPI) data using a 32-channel head coil (repetition time: 1.5 s; echo time: 0.73 ms; number of slices: 93; number of voxels: XX × YY; in-plane resolution: 1.5 × 1.5 mm; slice thickness: 1.5 mm; flip angle: 66°). Altogether, we collected 1840 volumes each, corresponding to thirty blocks of 8 trials each for a total of 240 trials in the main experimental task. Three participants saw their scan interrupted for technical problems and performed a lower number of blocks (21, 19 and 15 instead of 30, respectively). Anatomical images were acquired using a MPRAGE T1-weighted sequence that yielded images with a 0.7 × 0.7 × 0.7 mm resolution (160 slices; number of voxels: 256 × 256; repetition time: 8.2 ms; echo time: 3.7 ms).

## fMRI pre-processing

Pre-processing of our data was performed using the FMRIB’s Software Library (Functional MRI of the Brain, Oxford, UK) and included: head-related motion correction, slice-timing correction, highpass filtering (>100 s), and spatial smoothing (with a Gaussian kernel of 8 mm full-width at half maximum). To register our EPI image to standard space, we first transformed the EPI images into each individual’s high-resolution space with a linear six-parameter rigid body transformation. We then registered the image to standard space (Montreal Neurological Institute, MNI) using FMRIB’s Non-linear Image Registration Tool with a resolution warp of 8 mm.

## fMRI analyses

We performed whole-brain statistical analyses of functional data using a multilevel approach within the generalized linear model (GLM) framework, as implemented in FSL through the FEAT module: Y = Xβ + ε = β1X1 + β2X2 + … + βNXN + ε where Y is a T ×1(T time samples) column vector containing the times series data for a given voxel, and X is a T × N (N regressors) design matrix with columns representing each of the psychological regressors convolved with a hemodynamic response function specific for human brains. β is a N × 1 column vector of regression coefficients and ε a T × 1 column vector of residual error terms. Using this framework we initially performed a first-level fixed effects analysis to process each individual experimental run which were then combined in a second level mixed-effects analysis (FLAME 1 + 2) to combine data across subjects, treating participants as a random effect. For all analysis, we performed a cluster inference using a cluster-defining threshold of |Z| > 3.8 with a FWE-corrected threshold of P = 0.001. Applying this framework, we performed the GLMs highlighted below.

All GLMs included three unmodulated stick function regressors: (1) at the onset of the stimuli, (2) at the time of the choice (3) at the time the feedback appeared on the screen. Trials without pressure are defined as trials where pressure is negative according to equation 1, that is, when the goal had been already reached.

GLM 1: Included a parametric regressor modulated by the trial-by-trial pressure experienced by participants at the time of the offer one modulated by the reward and effort (separately for trials with or without pressure) chosen at time of choice.

GLM 2: Included parametric regressors for pressure, advancement and time left at time of choice and of feedback.

GLM 3: Included parametric regressors for time left at time of choice, for reward and effort (divided by trials with or without pressure) at time of choice, and pressure and pressure prediction error at time of feedback

GLM 4: Included parametric regressors for pressure at time of offer, signed difference in value (as computed by the model) between the higher and lower effort option at time of choice, and advancement and time left at time of feedback.

GLM 5: this GLM was identical to GLM 4 substituting the signed difference in value with an unsigned absolute difference in value between the two options.

GLM 6 and 7: identical to GLM 4 and 5 but with the value computed by a standard effort discounting model (where effort is a cost).

### ROIs

We drew masks of the cingulate regions based on anatomical maps^36^ of the cingulate cortex (with a threshold of 12) and of putamen following^85^.

### Extracting time-series data

We extracted time-series data from subject-specific ROIs for a psycho-physiological interaction (PPI) analysis (see below). We back-projected these clusters from standard space into each individual’s EPI (functional) space by applying the inverse transformations as estimated during registration (see fMRI pre-processing section). Finally, we computed average time-series data from all voxels in the back-projected clusters in each subject to serve as a physiological regressor in the PPI analysis. For all the cingulate clusters, time series were averaged between the two bilateral clusters.

### PPI analysis

Using the procedure described above, we extracted time-series data from individual clusters in 5 cingulate regions and two subdivisions of Putamen, which served as a seed region (that is, the physiological regressor—PHY) for a PPI analysis. This analysis was primarily designed to investigate the potential interaction of the area encoding pressure and pressure prediction error. We constructed our psychological (PSY) task regressor as a parametric regressor boxcar regressor with a step function in the interval between stimulus onset and response time or for 1,5 second following the presentation of the feedback.

## Data availability

The fMRI statistical maps from analyses and behavioural data generated in this study have been deposited in an Open Science Framework project. The raw fMRI data are protected and are not available due to data privacy laws. All data and code will be made available at the time of publication.

## Code availability

The code to generate the results and the figure, and the experimental paradigm, of this study is available in an Open Science Framework project. All code will be made available at the time of publication.

## Acknowledgements

MAJA was funded by a Biotechnology and Biological Sciences Research Council David Phillips Fellowship (BB/R010668/2) and a Jacobs Foundation Fellowship. We would like to thank Patricia Lockwood for comments on the project, as well as members of the Motivation and Social Neuroscience and Social Decision Neuroscience labs for their support.

## Authors contribution

M.A.P and M.A.J.A. designed the experiments. M.A.P and D.P collected the data for study 1 and 2. M.A.P and L.F. collected the data for study 3. M.A.P. analysed the data for study 1, M.A.P and D.P. analysed the data for study 2, M.A.P and L.F. analysed the data for study 3. M.A.P and M.A.J.A wrote the paper. All authors discussed the results and implications and commented on the manuscript at all stages.

