## supplementary material for "Neural and computational mechanisms of effort under the pressure of a deadline"

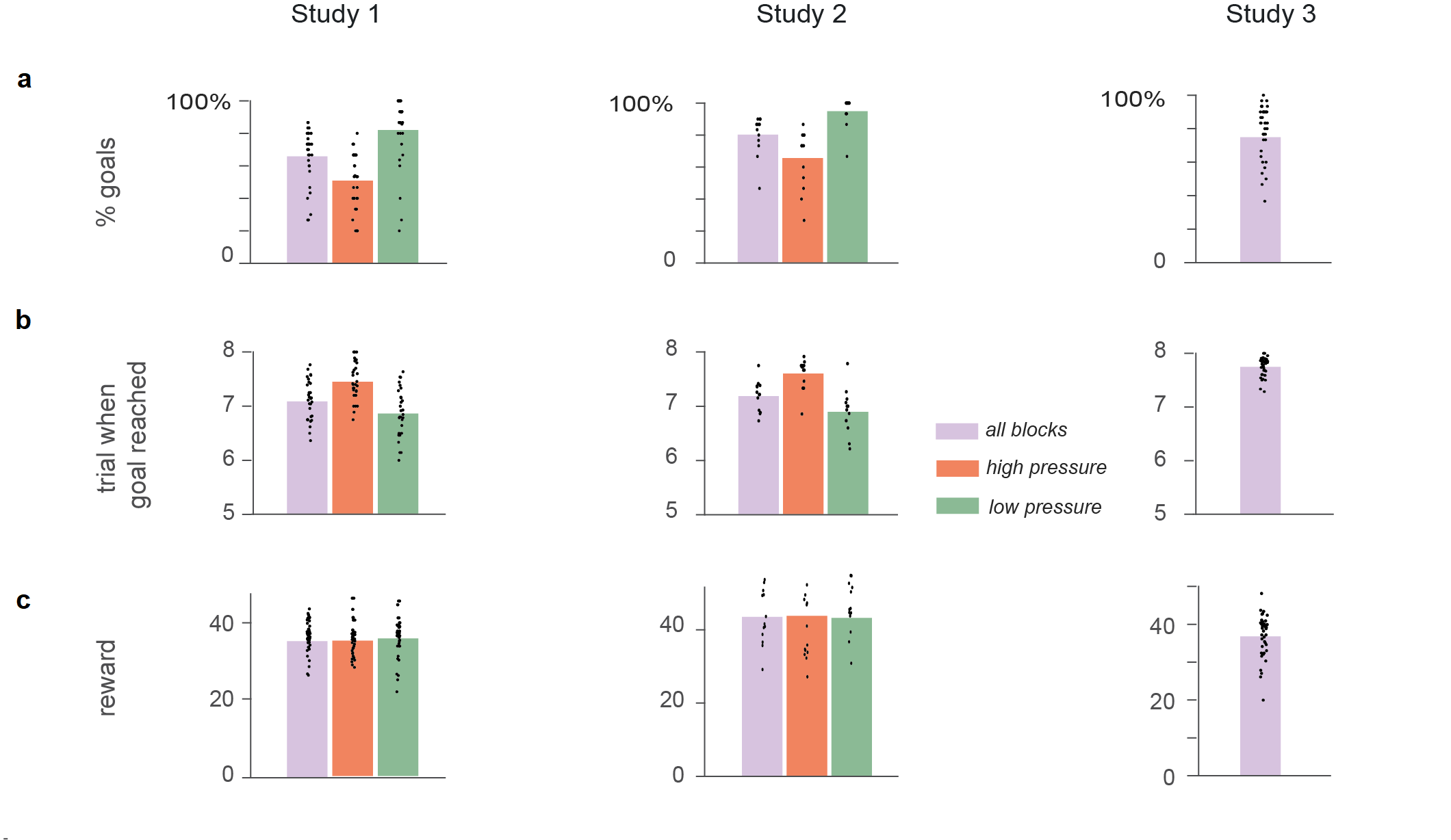

**Supplementary Figure 1.** *Performance in the task.* ***a****. Average percentage of goal reached across all, low high-pressure and low-pressure blocks in Study 1 (left), Study 2 centre) and Study 3 (right).* ***b****. Average number of trials before deadline in which the goal was reached across all, low high-pressure and low-pressure blocks (right panel) in Study 1 (left), Study 2 centre) and Study 3 (right).* ***c****. Average rewards collected across all, low high-pressure and low-pressure blocks (right panel) in Study 1 (left), Study 2 centre) and Study 3 (right).*

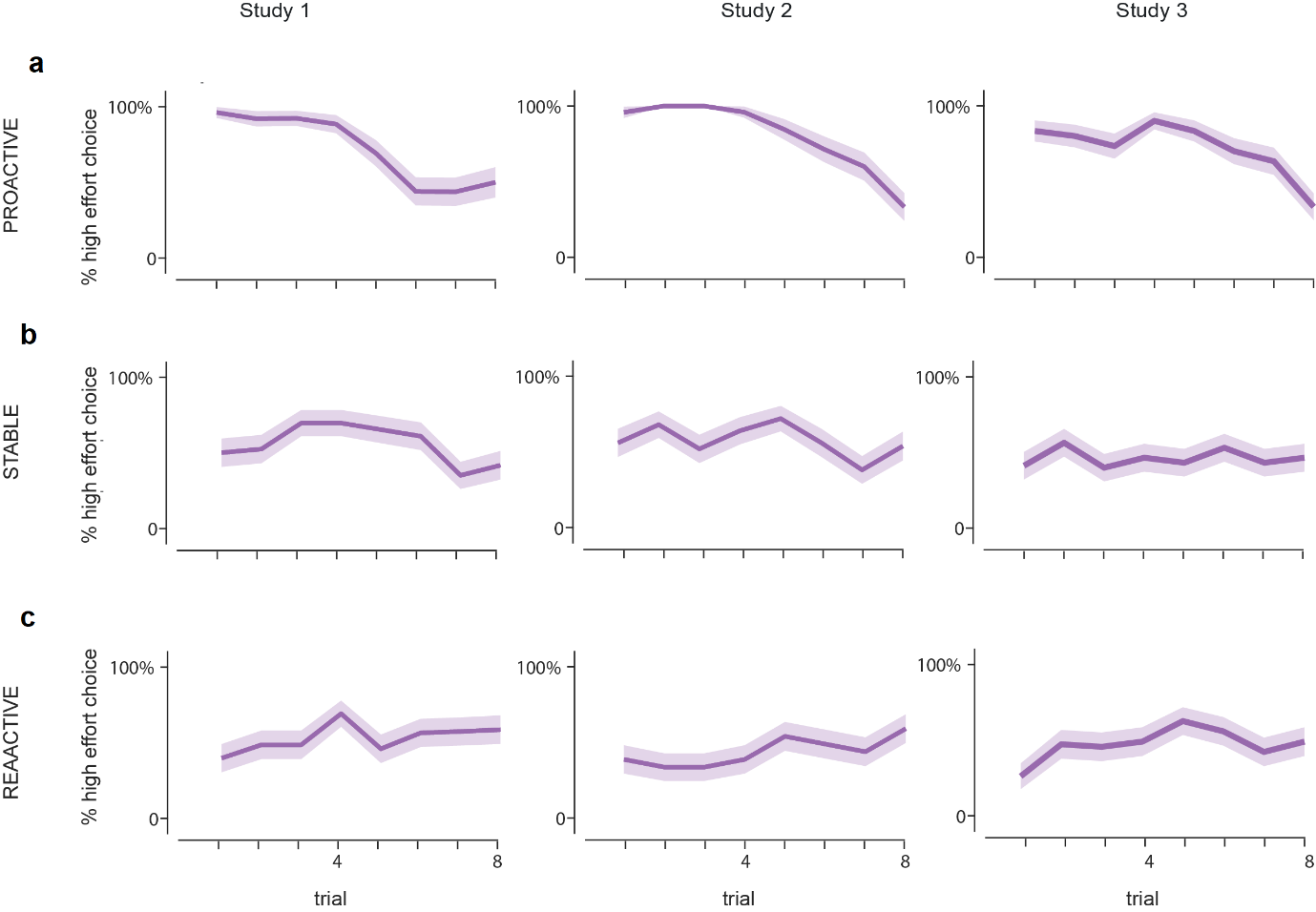
**Supplementary Figure 2.** *Examples of behaviour in the task.* ***a****. Individual subject average percentage of high effort choices across trials in Study 1 (left panel), Study 2 middle panel) and Study 3 (right panel). Participants could put in most effort at the beginning (proactive strategy, top row);* ***b****. distribute efforts equally across trials (stable strategy, middle row) or* ***c****. grow the likelihood to exert a high effort as time progresses (reactive strategy, bottom row).*

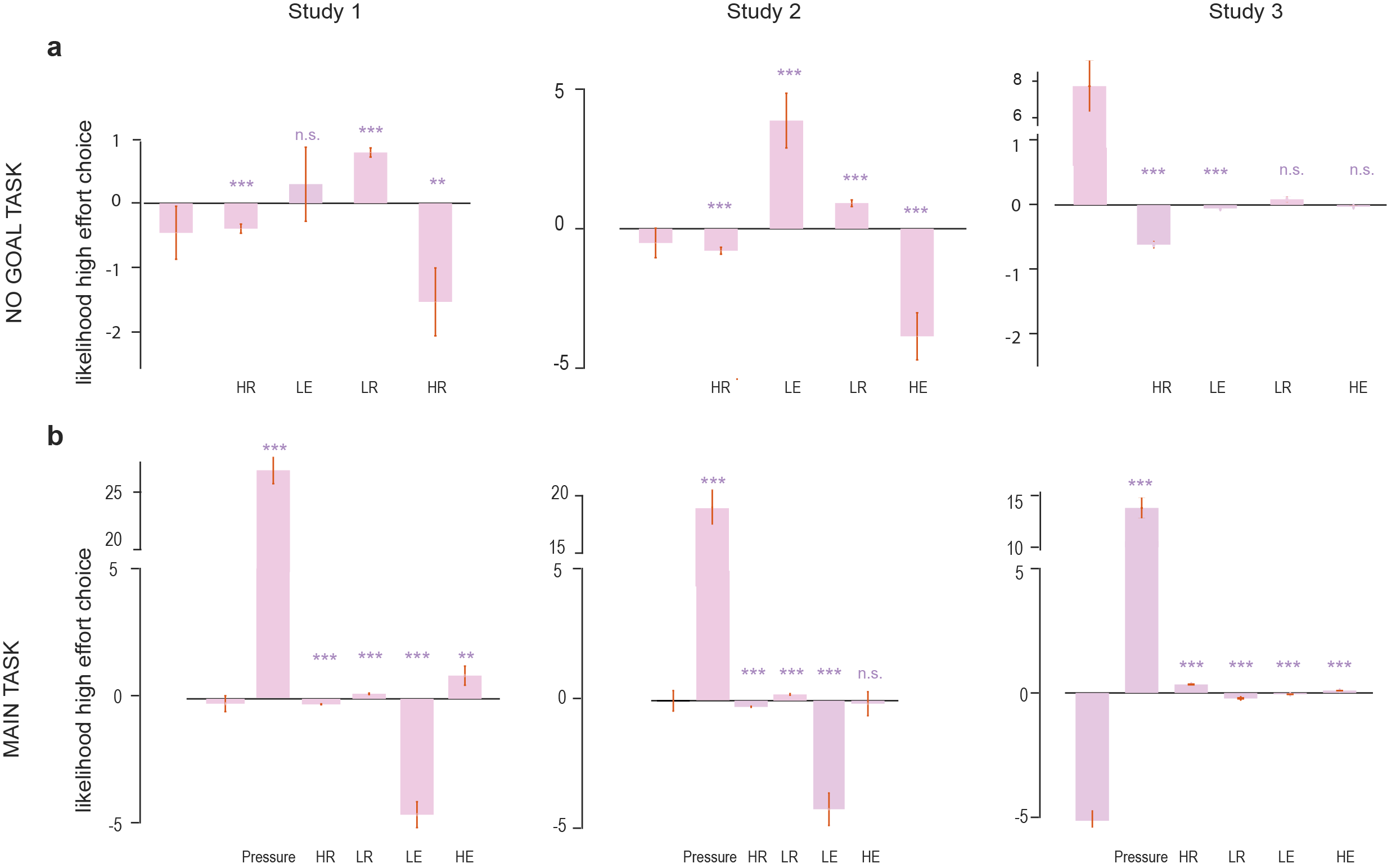

**Supplementary Figure 3.** *Predictors of behaviour in the task.* ***a****. A logistic regression predicting the likelihood of a High Effort Low Reward during the no-goal task in trials in Study 1 (left panel), Study 2 middle panel) and Study 3 (right panel).* ***b****. A logistic regression predicting the likelihood of a High Effort Low Reward during the main task in Study 1 (left panel), Study 2 middle panel) and Study 3 (right panel).*

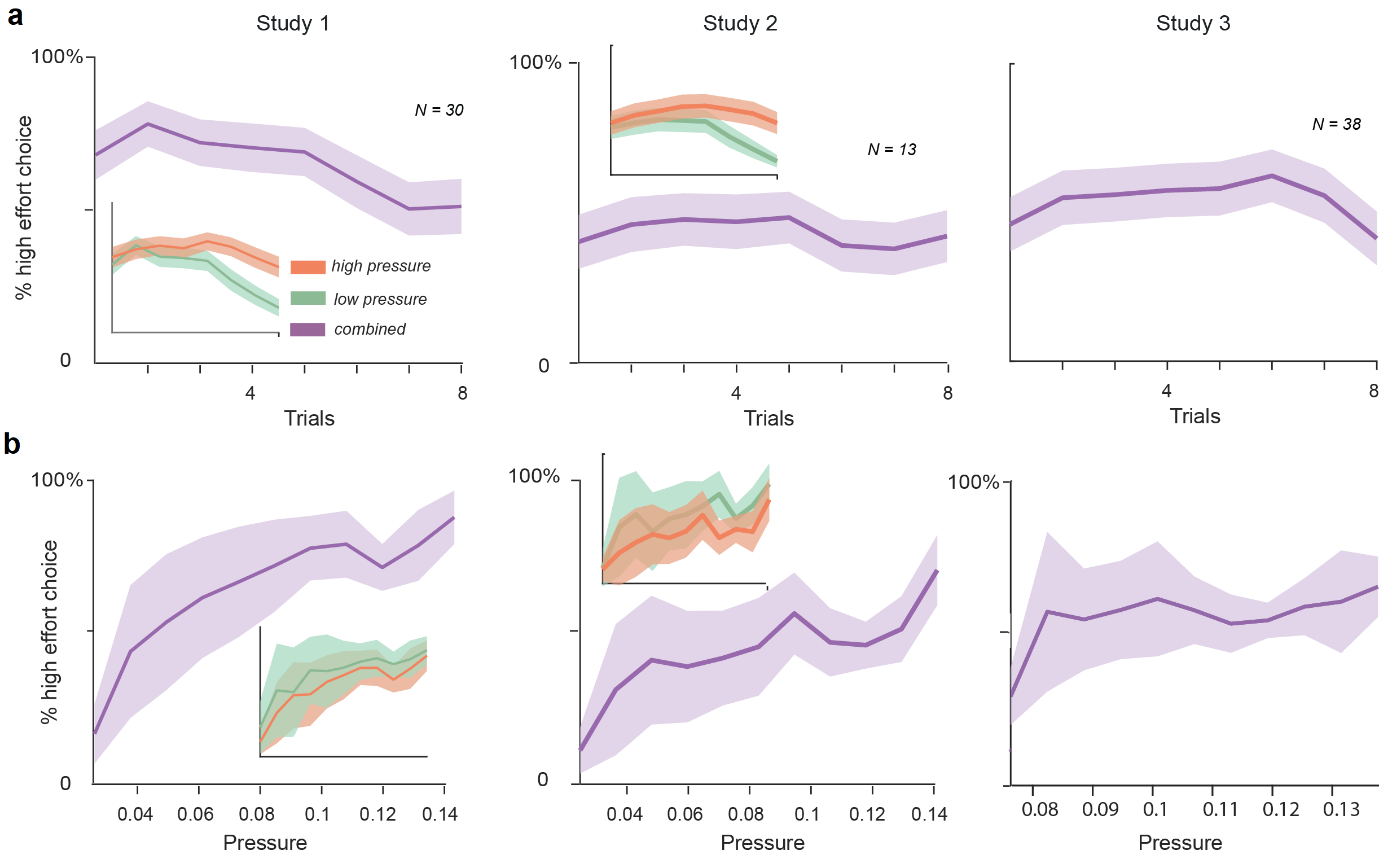

**Supplementary Figure 4.** *Behaviour in the task across studies.* ***a****. Average (across subjects and blocks) percentage of high effort choices across trials for study 1 (top row). In the inset, the same for high-pressure (red) and low-pressure (green) blocks. Average (across subjects and blocks) percentage of high effort choices across pressure values (bottom row). Pressure values where binned in 10 bins. In the inset, the same for high-pressure (red) and low-pressure (green) blocks.* ***b****. Same for study 2.* ***c****. same for study 3.*

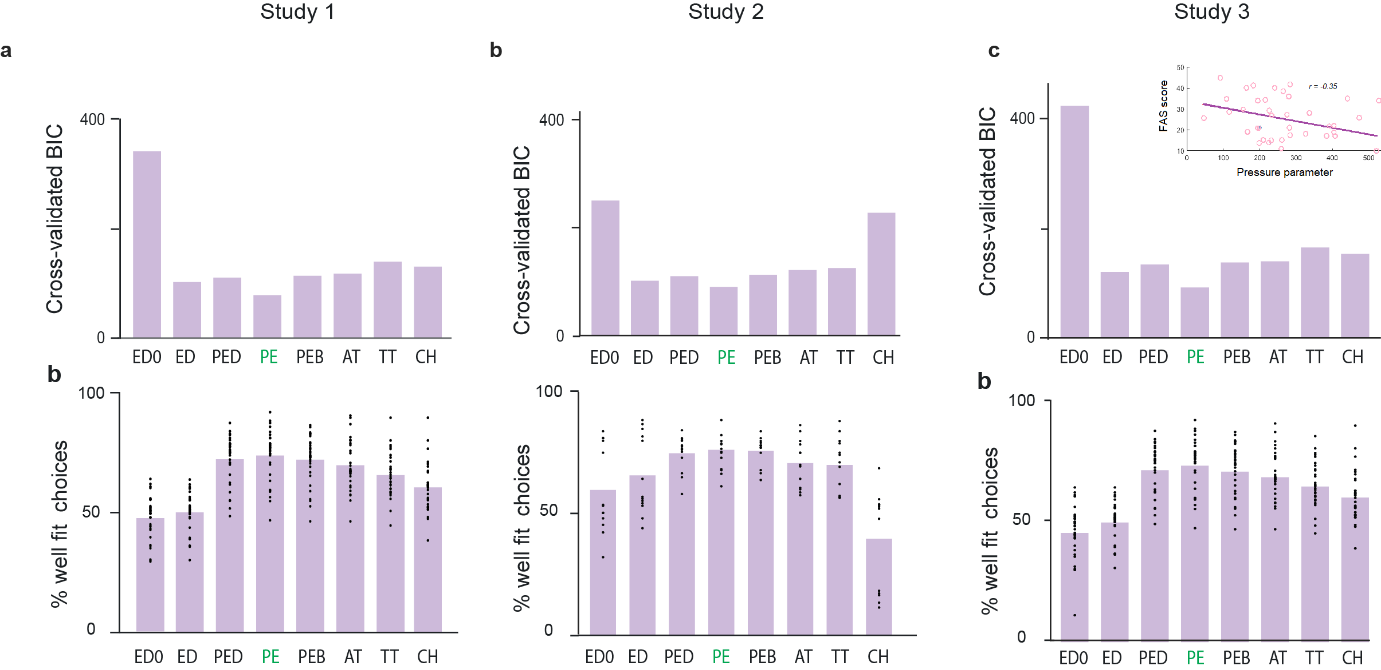

**Supplementary Figure 5** *Computational modelling.* ***a****. BIC values for different computational modelling of the evaluation and choice process for Study 1 (left), Study 2 (middle) and Study 3 (right). In all cases, the winning model is PE. In the inset, correlation between the pressure sensitivity parameter and the score in the Fatigue Assessment Scale, each point is a subject.* ***b*** *average percentage of correct choices across models for Study 1 (left), Study 2 (middle), Study 3 (right).*

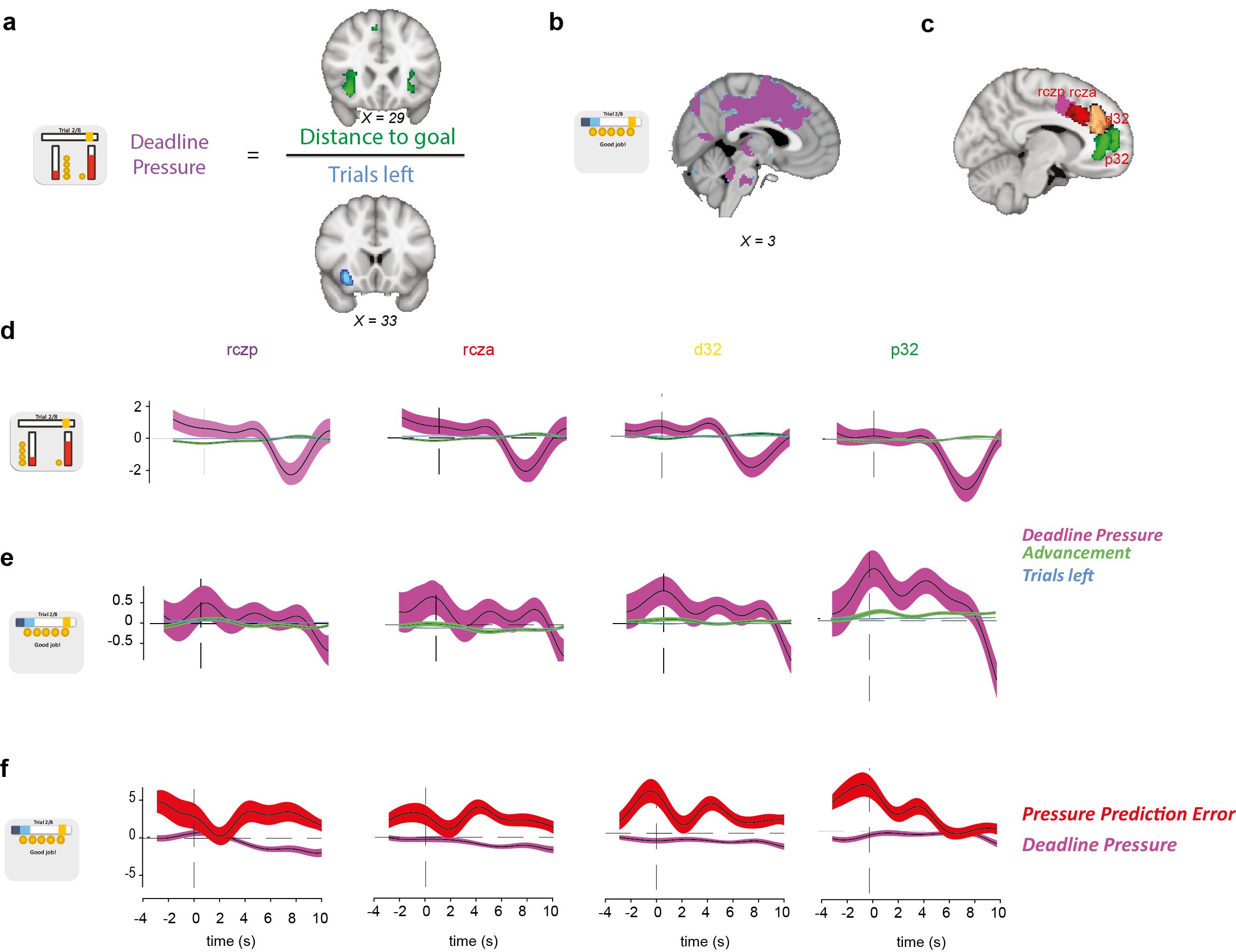

**Supplementary Figure 6.** *Neural representation in Middle and Anterior Cingulate Cortex.* ***a****. Representation of advancement and time left at time of offer in the right Insula.* ***b.*** *Representation of pressure at feedback time.* ***c****. Four anatomical masks for sections of the Middle and Anterior Cingulate Cortex.* ***d.*** *Time series of the strength of representation of Pressure (purple), Advancement (green), Time Left (cyan) at time of choice in voxels from the four anatomical masks.* ***e.*** *Time series of the strength of representation of Pressure (purple), Advancement (green), Time Left (cyan) at time of feedback in voxels from the four anatomical masks.* ***f****. Time series of the strength of representation of Pressure (purple) and Pressure prediction error (red) at time of feedback in voxels from the four anatomical masks.*

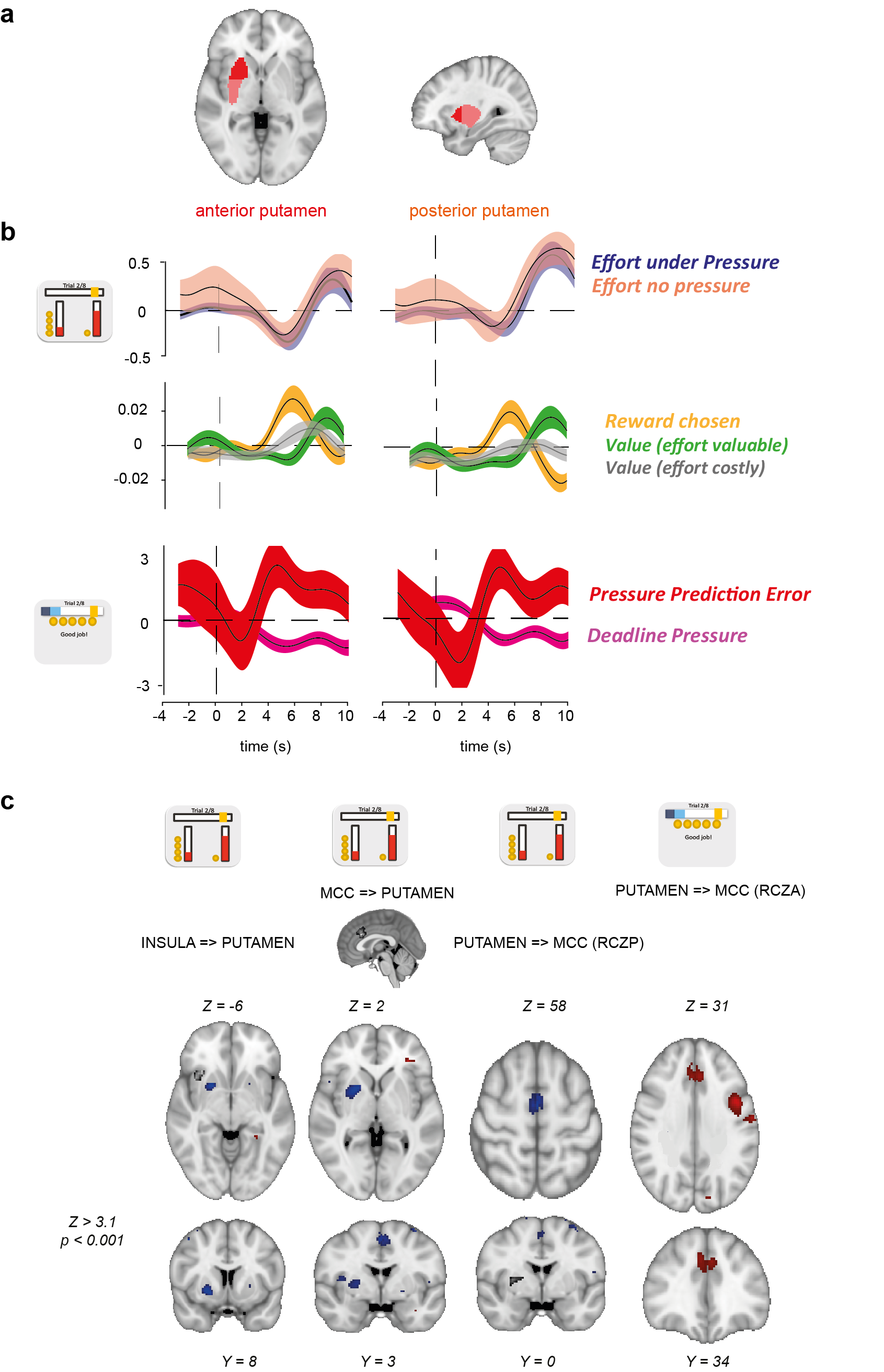

**Supplementary Figure 7.** *Neural representation in anterior and posterior Putamen.* ***a****. Anatomical subdivision of Putamen into anterior and posterior subregions.* ***b*** *Time series of the strength of representation of Effort, Reward, Value (at time of choice), Pressure and Pressure prediction error (at time of feedback) in voxels from the anterior and posterior Putamen anatomical ROIs.* ***c****. PPI analysis using clusters in Insula (first column) and Anterior Cingulate Cortex (second column) as a seed revealed an inverse coupling with the posterior putamen. PPI analysis using clusters in Putamen at time of choice (third column) or feedback (fourth column) as a seed revealed significant coupling with regions in RCZp and RCZa respectively.*

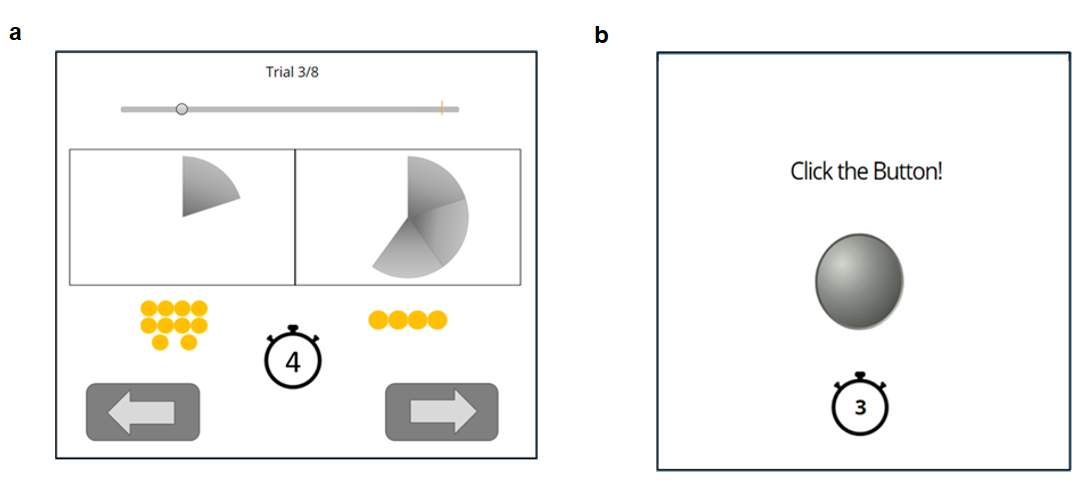

**Supplementary Figure 8.** *Description of the task for study 3, performed online.* ***a****. Participants need to reach a goal clicking a certain amount of times over 8 trials. The goal is represented by a little yellow line on the right hand side of the bar signalling the subject progress (top). In each trial subject are offered a choice between two different levels of efforts, represented by the slices in a pie chart, in exchange of two different amounts of reward, represented by the yellow coins. The choice was made by pressing either the left or right arrow buttons.* ***b****. After each decision, subjects are asked to click for the number of times associated with the chosen effort level.*

**Supplementary Table 1**

| **GLM 1** |  | **Peak MNI coordinates (mm)** | | |  |
| --- | --- | --- | --- | --- | --- |
| **Region** | **Hemisphere** | ***X*** | ***Y*** | ***Z*** | **Z Value (Peak)** |
| ***Pressure*** |  |  |  |  |  |
| Supramarginal Gyrus | R | 56 | -26 | 46 | -7.45 |
| Cingulate Gyrus (RCZp) | R | 2 | 2 | 36 | -5.78 |
| Putamen | R | 30 | 3 | -4 | -5.56 |
| Inferior Frontal Gyrus | R | 54 | 16 | 0 | -5.10 |
| Superior Frontal Gyrus | L | -30 | -2 | 68 | -5.10 |
| Lingual Gyrus | L | 24 | -52 | 2 | -4.33 |
| Lateral Occipital Cortex | R | 46 | -62 | 26 | -4.89 |
| Inferior Frontal Gyrus | R | 58 | 14 | 36 | -4.89 |
| ***Reward*** |  |  |  |  |  |
| Occipital Pole | R | 18 | -96 | 6 | 5.89 |
| Lingual Gyrus | L | -28 | -60 | -4 | 5.30 |
| ***Effort (under pressure)*** |  |  |  |  |  |
| Paracingulate Gyrus | L | -14 | 50 | 18 | -4.60 |
| Insula | L | -30 | 20 | -4 | -4.41 |
| Middle Frontal Gyrus | L | -44 | 26 | 32 | -4.19 |
| ***Effort (without pressure)*** |  |  |  |  |  |
| Middle Temporal Gyrus | L | -64 | -46 | -10 | 4.81 |
| Middle Frontal Gyrus | R | 36 | 34 | 36 | -6.95 |
| Insula | L/R | ±30 | 20 | -6 | -5.77 |
| Superior Parietal Lobule | L | -32 | -58 | 52 | -4.77 |
| Supramarginal Gyrus | L/R | 42 | -38 | 44 | -5.34 |
| Caudate | R | 10 | 12 | 8 | -4.89 |
| Frontal Pole | L | -30 | 46 | 0 | -4.15 |

| **GLM 2** |  | **Peak MNI coordinates (mm)** | | | | | |  | |
| --- | --- | --- | --- | --- | --- | --- | --- | --- | --- |
| **Region** | **Hemisphere** | ***X*** | | ***Y*** | | ***Z*** | | **Z Value (Peak)** | |
| ***Pressure (offer time)*** |  |  | |  | |  | |  | |
| Subcallosal Cortex | L | -1 | | 19 | | -5 | | -4.15 | |
| Lingual Gyrus | R | -8 | | -81 | | -6 | | -4.25 | |
| Occipital Pole | L | -14 | | -98 | | 14 | | 4.93 | |
| Middle Temporal Gyrus | R | 66 | | -48 | | -10 | | 4.95 | |
| ***Pressure (feedback time)*** |  |  | |  | |  | |  | |
| Fusiform Cortex | R/L | ±28 | | 52 | | -16 | | -4.97 | |
| Lingual Gyrus | L | -10 | | -64 | | -6 | | -4.30 | |
| Occipital Cortex | R | 26 | | -64 | | 46 | | -4.35 | |
| Inferior Temporal Gyrus | R | 46 | | -52 | | -14 | | -3.99 | |
| ***Advancement (offer time)*** |  |  | |  | |  | |  | |
| Insula | R/L | ±32 | | 26 | | -4 | | 5.93 | |
| Superior Frontal Gyrus | R | 4 | | 26 | | 48 | | 5.31 | |
| Angular Gyrus | R | 52 | | -48 | | 42 | | 4.92 | |
| Middle Frontal Gyrus | R | 40 | | 28 | | 42 | | 4.54 | |
| Supramarginal Gyrus | L | -42 | | -46 | | 48 | | 4.37 | |
| Frontal Pole | R | 30 | | 58 | | 0 | | 4.47 | |
| Cingulate Gyrus | R | 8 | | -38 | | 40 | | 4.39 | |
| Occipital Pole | L | -36 | | -90 | | 10 | | 4.18 | |
| Occipital Pole | L | -10 | | -94 | | 8 | | -4.31 | |
| ***Advancement (feedback time)*** |  |  | |  | |  | |  | |
| Brain Stem |  | 0 | | -38 | | -30 | | 4.78 | |
| Middle Temporal Gyrus | R/L | ±56 | | -54 | | 12 | | -6.69 | |
| Lobule Cortex | RL | ±4 | | -4 | | 62 | | -5.38 | |
| Precuneous Cortex | L | -6 | | -50 | | 54 | | -4.78 | |
| Lateral Occipital Cortex | R | 26 | | -70 | | 54 | | -4.70 | |
| Superior Parietal Lobule | L | -32 | | -52 | | 52 | | -4.42 | |
| Frontal Pole | R | 24 | | 58 | | 24 | | -5.21 | |
| ***Time Left (offer time)*** |  |  | |  | |  | |  | |
| Fusiform Gyrus | L | -18 | | -78 | | -8 | | 5.25 | |
| Lateral Occipital Cortex | R | 48 | | -74 | | 30 | | -5.19 | |
| Middle Temporal Gyrus | R | 66 | | -48 | | 0 | | -4.57 | |
| ***Time Left (feedback time)*** |  |  | |  | |  | |  | |
| Paracingulate Gyrus | R/L | ±8 | | 14 | | 44 | | -6.12 | |
| Postcentral Gyrus | R/L | ±42 | | -20 | | 42 | | -6.15 | |
| Superior Temporal Gyrus | R/L | ±58 | | -28 | | 0 | | -5.15 | |
| Occipital Pole | R/L | ±26 | | -90 | | -6 | | -5.83 | |
| Brain Stem | R/L | ±4 | | -22 | | -36 | | -5.36 | |
| Superior Parietal Lobule | L | -40 | | -40 | | 52 | | -4.95 | |
| Frontal Pole | L | -30 | | 42 | | 32 | | -4.38 | |
| Middle Frontal Gyrus | R | 32 | | 6 | | 62 | | -4.51 | |
| Caudate | R | 14 | | 14 | | 12 | | -4.35 | |
| Putamen | R | 29 | | -6 | | -2 | | -4.28 | |
| Inferior Frontal Gyrus | L | -44 | | 24 | | 20 | | -4.05 | |
| **GLM 3** |  | | **Peak MNI coordinates (mm)** | |  | |  | |  |
| **Region** | **Hemisphere** | | ***X*** | | ***Y*** | | ***Z*** | | **Z Value (Peak)** |
| ***Pressure*** |  | |  | |  | |  | |  |
| Brain Stem | R/L | | 0 | | -38 | | -26 | | 5.48 |
| Subcallosal Cortex | L | | -2 | | 0 | | -12 | | 4.45 |
| Thalamus | R | | 14 | | -22 | | 16 | | -5.95 |
| Paracingulate Gyrus | R/L | | ±3 | | 11 | | 48 | | -6.31 |
| Insula | R/L | | ±36 | | 14 | | -6.2 | | -6.31 |
| Postcentral Gyrus | R | | 52 | | -22 | | 48 | | -7.38 |
| ***Pressure Prediction Error*** |  | |  | |  | |  | |  |
| Fusiform Gyrus | R/L | | ±32 | | -84 | | -14 | | 5.44 |
| Lobule Cortex - RCZa | L | | -2 | | 2 | | 60 | | 5.17 |
| Middle Frontal Gyrus | L | | -40 | | 0 | | 54 | | 5.09 |
| Inferior Frontal Gyrus | R/L | | ±54 | | 14 | | 30 | | 4.83 |
| Postcentral Gyrus | R | | 36 | | -34 | | 54 | | 4.59 |
| Fusiform Gyrus | R | | 22 | | -84 | | 6 | | 5.39 |
| Posterior Cingulate Gyrus | R/L | | ±2 | | -40 | | 18 | | 4.67 |
| Postcentral Gyrus | L | | -58 | | -22 | | 22 | | 4.59 |
| Angular Gyrus | R | | 48 | | -50 | | 12 | | 4.50 |
| Middle Temporal Gyrus | R/L | | ±48 | | -34 | | -2 | | 4.37 |
| Lateral Occipital Cortex | R/L | | ±46 | | -66 | | 6 | | 4.61 |
| Brain Stem | R/L | | 0 | | -24 | | -28 | | 4.16 |
| Thalamus | L | | -8 | | -16 | | 12 | | 4.30 |
| Mid Cingulate Gyrus | R/L | | ±4 | | -22 | | 36 | | 4.16 |
| Precentral Gyrus | R | | 28 | | -10 | | 68 | | 4.35 |
| Putamen | L | | -20 | | 6 | | 2 | | 4.24 |
| ***Time Left*** |  | |  | |  | |  | |  |
| Fusiform Gyrus | L | | -28 | | -80 | | -6 | | 4.80 |
| Angular Gyrus | R | | 46 | | -52 | | 38 | | -5.54 |
| Insula | R | | 31 | | 19 | | -9 | | -5.59 |
| ***Reward*** |  | |  | |  | |  | |  |
| Frontal Pole | R | | 44 | | 46 | | 24 | | 4.78 |
| Posterior Cingulate Gyrus | R | | 10 | | -40 | | 32 | | 4.43 |
| Precentral Gyrus | R/L | | ±30 | | -8 | | 50 | | 4.29 |
| Fusiform Gyrus | L | | -20 | | -74 | | -6 | | 4.73 |
| Lateral Occipital Cortex | L | | -26 | | -82 | | 20 | | 4.83 |
| Superior Frontal Gyrus | R/L | | ±22 | | -2 | | 50 | | 4.71 |
| Occipital Pole | R | | 16 | | -94 | | 16 | | 5.56 |
| Postcentral Gyrus | L | | -42 | | -24 | | 54 | | 4.41 |
| Superior Parietal Lobule | L | | 36 | | -42 | | 42 | | 4.31 |
| Precentral Gyrus | L | | -44 | | 0 | | 38 | | 4.26 |
| ***Effort (under pressure)*** |  | |  | |  | |  | |  |
| Lingual Gyrus | R | | 12 | | -68 | | -10 | | 4.95 |
| Putamen | R | | 28 | | -2 | | -4 | | 4.72 |
| Paracingulate Gyrus – p32 | R | | 10 | | 50 | | 14 | | -4.27 |
| Paracingulate Gyrus | R | | 10 | | 36 | | 26 | | -4.19 |
| Subcallosal Cortex | R/L | | 0 | | 30 | | 0 | | -4.17 |
| ***Effort (without pressure)*** |  | |  | |  | |  | |  |
| Posterior Cingulate Gyrus |  | | 0 | | -46 | | 20 | | 5.55 |
| Middle Temporal Gyrus | R/L | | ±64 | | -28 | | -16 | | 5.13 |
| Frontal Pole | L | | -10 | | 66 | | 22 | | 4.69 |
| Frontal Pole | L | | -40 | | 38 | | -8 | | 4.49 |
| Postcentral Gyrus | L | | -8 | | -38 | | 58 | | 4.75 |
| Precentral Gyrus | L | | -40 | | -16 | | 40 | | 4.45 |
| Putamen | R/L | | ±6 | | -10 | | -2 | | 4.44 |
| Lingual Gyrus | R | | 10 | | -72 | | -6 | | 4.76 |
| Opercular Cortex | L | | -38 | | -20 | | 20 | | 4.75 |
| Anterior Cingulate Gyrus – d32 | R | | 6 | | 35 | | 22 | | -4.10 |
| Paracingulate Gyrus | R | | 10 | | 30 | | 40 | | -4.69 |
| Insula | R/L | | ±32 | | 20 | | -6 | | -4.2 |
| Middle Frontal Gyrus | R | | 34 | | 34 | | 28 | | -4.34 |
| **GLM 4-7** |  | | **Peak MNI coordinates (mm)** | |  | |  | |  |
| **Region** | **Hemisphere** | | ***X*** | | ***Y*** | | ***Z*** | | **Z Value (Peak)** |
| ***Value Difference (signed)*** |  | |  | |  | |  | |  |
| Lobule Cortex | R | | -4 | | -10 | | 54 | | 4.19 |
| Lingual Gyrus | R/L | | 0 | | -66 | | 0 | | 4.76 |
| Anterior Cingulate Gyrus | R/L | | ±4 | | 36 | | 16 | | -4.11 |
| Paracingulate Gyrus | L | | -10 | | 52 | | 20 | | -4.13 |
| ***Value Difference (R-L)*** |  | |  | |  | |  | |  |
| Precentral Gyrus | L | | -32 | | -26 | | 70 | | 4.46 |
| Paracingulate Gyrus | R/L | | ±4 | | 36 | | 26 | | -5.31 |
| Inferior Frontal Gyrus | R | | 48 | | 24 | | 10 | | -5.89 |
| ***Value Difference (signed) - effort discounting model*** |  | |  | |  | |  | |  |
| Occipital Pole | R/L | | ±12 | | -90 | | -8 | | 6.08 |
| Postcentral Gyrus | L | | -40 | | -24 | | 54 | | 4.36 |
| Supramarginal Gyrus | L | | -48 | | -42 | | 52 | | 4.52 |
| Superior Frontal Gyrus | L | | -24 | | -6 | | 52 | | 4.29 |
| ***Value Difference (R-L) - effort discounting model*** |  | |  | |  | |  | |  |
| Anterior Cingulate Gyrus | L | | 0 | | 32 | | -2 | | 4.2 |
| Brain Stem | R/L | | ±2 | | -38 | | -26 | | 4.13 |
| Lingual Gyrus | L | | -18 | | -74 | | -6 | | 3.98 |
| Occipital Pole | L | | -10 | | -98 | | 10 | | -4.49 |

**Supplementary Table 1.** BOLD activations in the parametric regressors of our GLMs. Complete list of activations correlating negatively or positively with the single-trial variability. (GLM; mixed effects, |Z| *>* 3.8, corrected). MNI, Montreal Neurological Institute; L, left hemisphere; R, right hemisphere. Coordinates at peak, for bilateral clusters, coordinates of the highest peak.
